# Coherent Cross-modal Generation of Synthetic Biomedical Data to Advance Multimodal Precision Medicine

**DOI:** 10.1101/2025.08.22.671728

**Authors:** Raffaele Marchesi, Nicolò Lazzaro, Walter Endrizzi, Gianluca Leonardi, Matteo Pozzi, Flavio Ragni, Stefano Bovo, Monica Moroni, Venet Osmani, Giuseppe Jurman

## Abstract

Integration of multimodal, multi-omics data is critical for advancing precision medicine, yet its application is frequently limited by incomplete datasets where one or more modalities are missing. To address this challenge, we developed a generative framework capable of synthesizing any missing modality from an arbitrary subset of available modalities. We introduce Coherent Denoising, a novel ensemble-based generative diffusion method that aggregates predictions from multiple specialized, single-condition models and enforces consensus during the sampling process. We compare this approach against a multi-condition, generative model that uses a flexible masking strategy to handle arbitrary subsets of inputs. The results show that our architectures successfully generate high-fidelity data that preserve the complex biological signals required for downstream tasks. We demonstrate that the generated synthetic data can be used to maintain the performance of predictive models on incomplete patient profiles and can leverage counterfactual analysis to guide the prioritization of diagnostic tests. We validated the framework’s efficacy on a large-scale multimodal, multi-omics cohort from The Cancer Genome Atlas (TCGA) of over 10,000 samples spanning across 20 tumor types, using data modalities such as copy-number alterations (CNA), transcriptomics (RNA-Seq), proteomics (RPPA), and histopathology (WSI). This work establishes a robust and flexible generative framework to address sparsity in multimodal datasets, providing a key step toward improving precision oncology.

**Author Summary:** To make precision medicine a reality, doctors need to understand a patient’s status from many angles, using different data types like genetic information (omics) and tissue slide images (histopathology). The problem is that most patient records are incomplete, with one or more of these data types missing, which can limit the effectiveness of powerful predictive tools. We have built a generative AI system designed to learn the complex biological patterns that connect all these different data types. By looking at the patient data that is available, our system can then generate a realistic, synthetic version of any missing piece. We developed a novel method called Coherent Denoising to do this, which is flexible and helps protect patient privacy. We validated this approach on a large dataset of over 10,000 cancer patient profiles. We show that our AI-generated data is high-fidelity and can successfully complete these sparse patient profiles, allowing AI models for crucial tasks like cancer staging and survival prediction to work at their best even with incomplete patient data. We also demonstrate how this tool can be used to evaluate the potential impact of new tests, helping to prioritize which expensive diagnostic tests would be most beneficial for a patient.

## Introduction

Biomedical research increasingly relies on the integration of heterogeneous data to characterize complex biological systems. Traditional approaches often isolated specific modalities, treating them as independent analytical domains [1]. While this sectoral approach has led to substantial progress in many individual disciplines, it limits the ability to capture cross-scale dependencies and emergent patterns that arise from the interaction of multiple biological processes.

In recent years, a growing need has emerged to move beyond a fragmented view and adopt a more holistic, multidisciplinary approach. This involves aggregating, integrating, and analysing diverse biological scales and data modalities that comprehensively capture biomedical features across large patient populations [2, 3, 4, 5, 6]. Several studies have shown that this multimodal method can outperform single-modality approaches in specific tasks [7, 8]. One strategy for this integration is to operate on a unified embedding level [9]. This paradigm has been advanced by the emergence of large-scale foundation models, which can be pre-trained on vast datasets to learn powerful, dense representations of complex data [10]. For example, vision transformers like Titan [11] can distil the rich information from gigapixel whole-slide images into a single feature vector. Once each data type is converted into this common embedding format, they are typically fused within a shared latent space to enable downstream prediction [12].

This evolving landscape encouraged the proliferation of initiatives such as the The Cancer Genome Atlas Program (TCGA), the National Cancer Database (NCDB), the UK Biobank, which systematically collect and harmonize multi-omics, imaging, and clinical data across diverse populations to support biomarker discovery, disease modeling, and precision medicine applications [13, 14]. However, despite the advancements enabled by these large-scale projects, translating integrative models into clinical settings remains challenging [15]. Many patients datasets are inherently incomplete, often because certain modalities are prohibitively expensive, technically challenging to acquire routinely [16, 17], or simply unavailable in low-resource settings and specialized centers with limited access to comprehensive molecular diagnostics or advanced imaging equipment [18, 19]. Generative artificial intelligence (GenAI) is emerging as a powerful tool in this field, offering methods to model the complex distribution of medical data. Primary applications include data augmentation to expand or balance limited datasets, the generation of fully synthetic yet biologically plausible data to facilitate research while preserving patient privacy, and the synthesis of missing modalities to address data sparsity. Early frameworks based on Generative Adversarial Networks (GANs) [20, 21, 22, 23] and Variational Autoencoders (VAEs) [24] showed success in unimodal synthesis. For example, OmicsGAN [25] was used to enrich two omics modalities (e.g., mRNA and microRNA) using adversarial learning, integrating prior knowledge of molecular interaction networks to guide realistic data generation. Similarly, MG-GAN [26] was able to produce synthetic gene expression data for traing set augmentation in downtream tasks. In a complementary approach, RNA-GAN [27] demonstrates the potential of cross-modal learning generating synthetic whole-slide images (WSI) tiles, conditioned on RNA-sequencing profiles. More advanced architectures also incorporate multi-conditioning strategies, such as CLUE [28], a VAE-based single-cell data reconstruction tool. However, clinical translation of GANs and VAEs has been challenged by training instability, lower-fidelity outputs, and difficulties in extending them to flexible, conditional multimodal generation [29, 30, 31]. More recently, Denoising Diffusion Probabilistic Models (DDPMs) have emerged as a new state-of-the-art, offering stable training and high-quality sample generation [32, 33]. Their effectiveness is well-established for single-modality tasks [34, 35], particularly imaging [36, 37, 38]. For instance, MCAD [39] applies multi-conditional strategies to PET image reconstruction, while stDiffusion [40] enables the generation of spatial transcriptomics data. Nevertheless, at the current state-of-the-art, application of GenAI to the complex, any-to-any conditional synthesis of multimodal healthcare data remains a critical and underexplored research area [41].

To address these challenges in multimodal learning, we introduce a unified, cross-omics, cross-modal generative AI framework designed to synthesize missing biomedical data from any combination of available modalities. We demonstrate its capabilities on a large-scale, multimodal, multi-omics cancer dataset comprising copy-number alterations (CNA), transcriptomics (RNA-seq), proteomics (RPPA) or histopathology embeddings (WSI). Our primary contributions include: **(1)** ensemble generation via Coherent Denoising, a novel and highly scalable late-fusion ensemble method that aggregates predictions from multiple single-condition diffusion models, enforcing consensus during the sampling process; **(2)** a comparison of this approach against a state-of-the-art multi-condition diffusion model that seamlessly handles arbitrary subsets of input modalities with a flexible masking strategy; **(3)** a comprehensive validation on a pancancer cohort of over 10,000 primary tumors spanning 20 cancer types, with four data modalities, each encoded in a low-dimensional latent space; demonstrating that our methods not only reconstruct data with high fidelity but also preserve the critical biological signals required for downstream tasks, including tumor type classification, stage prediction, and survival analysis; highlighting the ensemble’s privacy-preserving advantages; **(4)** a showcase of the translational utility of our approach through two key applications: enhancing the performance of multimodal machine learning models in the face of missing data at inference time, and a novel counterfactual analysis to guide the strategic prioritization of data acquisition for diagnostics. Together, these results establish cross-modal generation as a robust tool to work with sparse patient profiles, with promising applications in precision medicine, in silico trials, and resource-constrained diagnostic workflows.

## Results

We utilized a large pan-cancer cohort from TCGA, comprising 10,098 samples with four distinct data modalities: CNA, RNA-Seq, RPPA, and WSI embeddings. To create a harmonized data representation suitable for multimodal learning, we first encoded each omics modality into a dense, 32-dimensional latent space using modality-specific autoencoders. We then developed and benchmarked two diffusion-based generative frameworks - a multi-condition model and our novel ensemble generation via Coherent Denoising - to synthesize any missing modality by conditioning on any combination of available ones. The full methodological process is described in the Methods section.

The efficacy of our generative framework was evaluated through a series of experiments on a held-out test set of 1,350 pan-cancer samples. The evaluation first assesses the fidelity of reconstruction, beginning with a qualitative analysis of the global data manifold using Uniform Manifold Approximation and Projection (UMAP), followed by a quantitative measurement of the performance for each modality using the coefficient of determination. Subsequently, we use a downstream classification task as a functional test of generation quality, evaluating if the synthetic data preserves the predictive signals required by single-modality classifiers. Finally, we demonstrate two practical applications: the use of generative data completion to enable high performance of a multimodal predictive model faced with incomplete test data and a counterfactual inference approach to guide the prioritization of data acquisition. We also note an inherent privacy-preserving advantage of the ensemble approach, which unlike a monolithic multi-condition model proves robust against unconditional generation aimed at reconstructing the training data manifold.

### Multimodal Distribution Fidelity

As a first step, we investigated whether our generative frameworks could preserve the global structure of the multimodal data manifold. This analysis qualitatively evaluates the preservation of large-scale biological patterns, namely the distinct clustering between cancer types, evident in the original data. To this end, we performed a qualitative analysis using the UMAP [42]. While we acknowledge that UMAP may not always preserve global structure accurately [43], we use to assess the visual correspondence between the real and generated data manifolds, rather than as a definitive clustering tool.

We compared the UMAP embedding of the ground-truth test set against embeddings from two fully reconstructed test sets. These were generated by our primary methods: the multi-condition model and the Coherent Denoising ensemble model. For this analysis, a complete, synthetic profile for each test sample was assembled by iteratively reconstructing each of the four omics and modalities (CNA, RNA-Seq, RPPA, WSI) while conditioning on the real, ground-truth versions of the other three. The four resulting synthetic modalities for each patient were then concatenated to form a final reconstructed embedding vector.

Fig 1 presents the side-by-side comparison of these UMAP projections. The ground-truth data (Fig 1, Left) reveals a well-defined structure where samples form distinct clusters that correspond closely to their cancer type of origin. This underlying organization confirms that the concatenated multimodal embeddings effectively capture strong, tissue-specific biological signatures. The reconstructed data from both the multi-condition model (Fig 1, Middle) and the Coherent Denoising method (Fig 1, Right) demonstrate a high degree of qualitative similarity to the ground-truth manifold. Both generative approaches successfully reproduce the primary clusters with respect to their relative positions, shapes, and degree of separation. The UMAP visualizations of each individual modality, presenting patterns similar to those in Fig 1, are shown in S1 Appendix.

**Fig 1.**
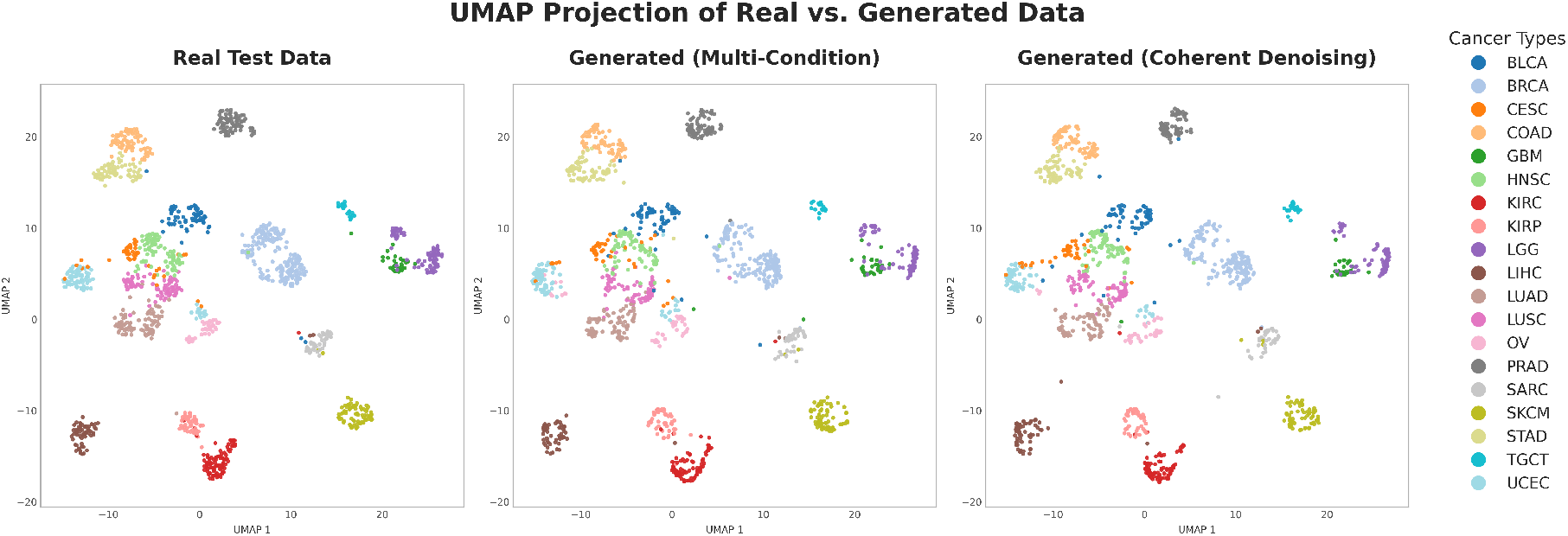
Qualitative comparison of real and generated data manifolds. UMAP projections of multimodal embeddings from the test set. The distribution of ground-truth data (Left) is compared with data reconstructed by the multi-condition model (Middle) and the Coherent Denoising ensemble (Right). Each point is a patient sample, colored by cancer type. Both generative approaches successfully capture the global topology and distinct cancer-type clustering of the original data.

Visual inspection indicates that the overall topology of the data space is well-preserved by both methods. Crucially, our framework achieves this by generating each target modality conditioned on all the other available modalities, rather than performing a single, joint multimodal generation. The generated samples are not randomly distributed but are consistent with the complex biological structure of the original data, successfully synthesizing the multimodal signatures that differentiate major cancer types. This qualitative validation provides good initial evidence that our framework can generate multimodal patient profiles that are both diverse and biologically plausible, a critical prerequisite for their use in downstream applications.

### Reconstruction Fidelity

Following the qualitative validation, we performed a quantitative analysis to assess the fidelity of our generative frameworks’ reconstructions. We evaluated the performance of both the multi-condition model and the ensemble generation via Coherent Denoising across all possible combinations of conditioning modalities. The coefficient of determination (*R*^2^) was used as the primary metric, where a value of 1.0 indicates perfect reconstruction and a negative value indicates poorer performance than a baseline model simply predicting the mean. For each combination of model and conditioning input, the entire test set was reconstructed 10 independent times. We then calculated *R*^2^ for each of the 10 runs. To assess model consistency, we also measured the output variance by calculating the variance across the 10 generated versions for each individual test sample, averaging this per-sample variance across the entire test set, and expressing it as a percentage of the real data variance. The full results for both metrics are reported in Table 1.

**Table 1.** Reconstruction accuracy (*R*^2^) and output variance by target modality and generative method. Each row shows the mean *R*^2^± standard deviation from *10 independent generation runs*. Columns indicate the generative method used: single-condition models, multi-condition models, and Coherent Denoising. The best *R*^2^ for each target modality is highlighted in bold. Values in parentheses show the average variance of the generated samples as a percentage of the total variance from the real data; lower percentages indicate less output variability, indicating confidence of the model in the sample generation.

| Target modality | Single-condition |  |  |  | Multi-condition | Coherent Denoising |
| --- | --- | --- | --- | --- | --- | --- |
|  | cna | rnaseq | rppa | wsi |  |  |
| cna | — | -0.094 ± 0.011 (16.0%) | -0.160 ± 0.012 (12.3%) | -0.172 ± 0.011 (12.8%) | -0.064 ± 0.011 (10.4%) | **0.063 ± 0.016 (13.1%)** |
| rnaseq | 0.318 ± 0.012 (12.6%) | — | 0.770 ± 0.001 (0.8%) | 0.702 ± 0.001 (1.0%) | **0.787 ± 0.001 (0.4%)** | 0.746 ± 0.001 (0.8%) |
| rppa | 0.169 ± 0.016 (12.1%) | 0.605 ± 0.001 (1.1%) | — | 0.523 ± 0.002 (1.4%) | **0.620 ± 0.001 (0.8%)** | 0.558 ± 0.001 (1.1%) |
| wsi | -0.051 ± 0.006 (10.6%) | 0.361 ± 0.003 (3.0%) | 0.358 ± 0.001 (2.1%) | — | 0.392 ± 0.002 (1.2%) | **0.439 ± 0.002 (2.1%)** |

The results are summarized in Table 1. Detailed metrics, complete conditional combinations, and a stratified analysis by individual cancer type are provided in S2 Appendix. The analysis revealed several key findings.

First, the reconstructibility of the four data types varied significantly. RNA-Seq was the most successfully reconstructed modality, achieving a mean *R*^2^ of 0.79 with the multi-condition model. This high fidelity was accompanied by extremely low output variance (just 0.4% of the real data’s variance), indicating low model’s uncertainty. RPPA and WSI embeddings were also reconstructed effectively, reaching maximum mean *R*^2^ scores of 0.62 and 0.44, respectively, with similarly low output variance (0.8% and 2.1%). In contrast, CNA data proved exceptionally challenging to generate from other data types. The best-performing model for CNA reconstruction achieved a mean *R*^2^ of only 0.06, with most model combinations yielding negative scores. This suggests that the information preserved within the highly compressed CNA embedding is largely uncorrelated with the latent representations of the other modalities. The poor reconstruction of the challenging CNA modality was marked by substantially higher output variance (up to 16.0%). This shows that the generative process is able to explicitly model its high uncertainty in light of insufficient information.

Second, reconstruction performance generally improved when more conditioning modalities were provided. For instance, generating WSI data from RNA-Seq alone yielded a mean *R*^2^ of 0.36, which increased to 0.43 when both RNA-Seq and RPPA data were used as inputs for the Coherent Denoising model. While performance gains saturate for high-fidelity modalities like RNA-Seq, the framework is designed to robustly handle any arbitrary subset of inputs, which is essential for typical sparse data scenarios (S2 Appendix).

Finally, there is performance trade-off between Coherent Denoising ensemble and multi-condition model contingent on the target omic or modality. The multi-condition model achieved the highest fidelity for the most predictable targets, RNA-Seq (*R*^2^=0.79) and RPPA (*R*^2^=0.62). Conversely, the Coherent Denoising ensemble demonstrated superior performance for the more challenging targets, outperforming the multi-condition model for WSI (*R*^2^=0.44 vs. 0.39) and proving more effective in modeling high-uncertainty (CNA data *R*^2^=0.06 vs.-0.06).

### Preservation of Predictive Signals in Generated Data

To evaluate generation quality beyond reconstruction metrics, we designed an experiment to measure whether synthetic data retain the complex biological signals necessary for downstream predictive tasks. This experiment serves as a functional test of generation fidelity. The objective is not to compare which input is more predictive, but rather to verify if the same classifier, trained only on real data, can achieve equivalent performance when tested on the synthetic data. We trained Random Forest classifiers [44] with 500 estimators (trees) on the real training set to perform two distinct tasks: 20-class tumor type prediction and 4-class cancer stage prediction. Separate classifiers were trained for each of the four data modalities, using only the real data from the training set. Random Forests were chosen for their robustness to high-dimensional data, ability to capture non-linear relationships, and consistently strong performance across diverse biological datasets [45, 46, 47]. To ensure stable results and robust evaluation, for each generative method and target modality, a synthetic test set was generated 10 times, each using a different random seed for the generative process. The performance of the already trained classifier was then evaluated on each of these 10 generated test sets.

The quality of the generated data was then assessed by comparing the classifier’s performance on the real test data with its performance on synthetic data. This synthetic data was generated by our multi-condition and Coherent Denoising models, both conditioned on the remaining three real modalities. To account for class imbalance, Fig 2 shows the mean macro F1-score and standard deviation across the 10 experimental replicates. Full detailed metrics, including balanced accuracy, are available in S3 Appendix. For reference, a baseline performance of a random classifier would be expected to achieve a macro F1-score of 0.05 for tumor type prediction and 0.25 for stage prediction.

**Fig 2.**
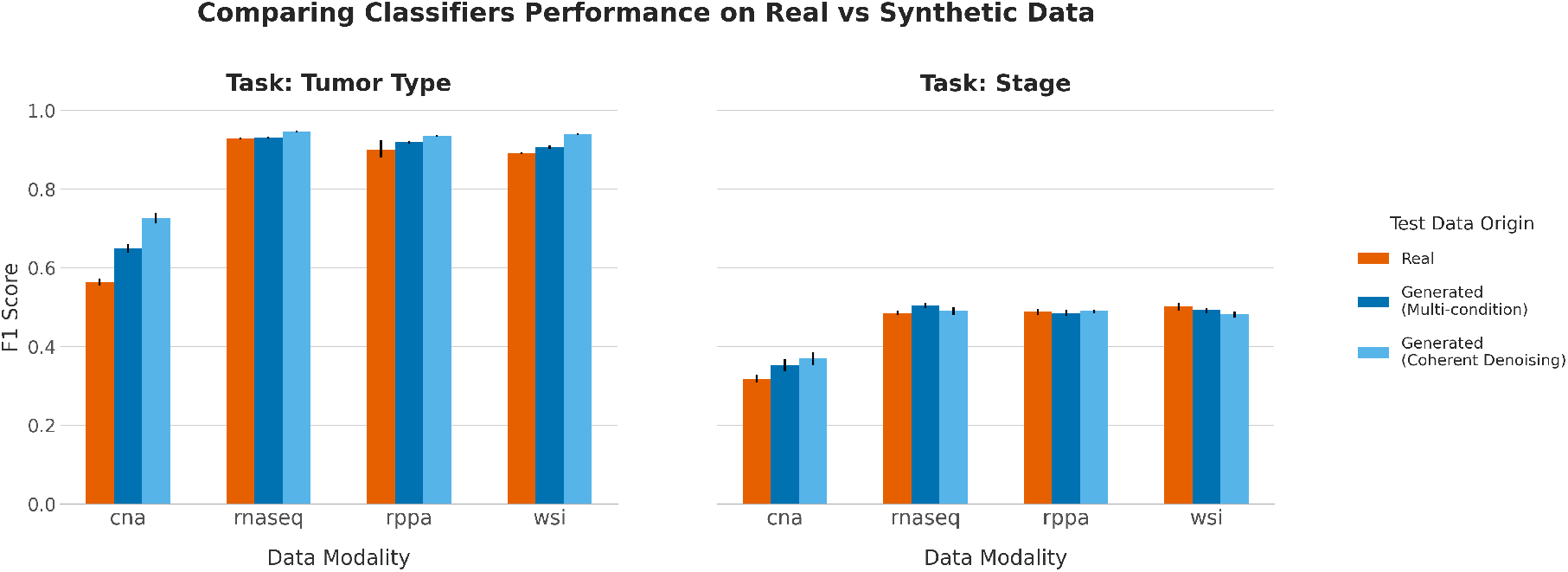
Classification performance on real versus synthetic data. Macro F1 scores for Random Forest classifiers trained on single real modalities and tested on either real data (orange), data from the multi-condition model (dark blue), or data from the Coherent Denoising model (light blue). The left panel shows results for the 20-class tumor type prediction task, and the right panel shows results for the 4-class stage prediction task. Error bars represent the standard deviation across 10 independent experimental runs.

For modalities already rich in predictive information, such as RNA-Seq, RPPA, and WSI, the classifiers’ performance on synthetically generated data was nearly identical to their performance on real data. In the tumor type prediction task (Fig 2, Left), a classifier using real RNA-Seq data achieved a F1-score of 0.94. The same classifier, when tested on RNA-Seq data generated by our Coherent Denoising model, achieved a F1-score of 0.95. The only instance of a minor, but statistically significant performance decrease was observed for WSI data generated by the Coherent Denoising model for the stage prediction task: from F1-score of 0.50 to 0.48 (see S6 Appendix, Section 1 for all statistical tests). This parity in performance indicates that the generative process successfully captures and reconstructs the features within these modalities that are most relevant for prediction.

A notable divergence in performance was observed with CNA data, a modality with weaker intrinsic predictive power compared to RNA-Seq, RPPA, or WSI. As depicted in Fig 2 (Left), classifiers tested on synthetic CNA data yielded significantly higher prediction scores than when evaluated on real CNA data. For instance, in the tumor type prediction task, the mean balanced accuracy for CNA data increased from 0.565 on real data to 0.717 on data generated by the Coherent Denoising model (see S3 Appendix). This suggests that the genomic alterations of a patient’s CNA are not fully reconstructible from other modalities (consistent with the low *R*^2^ scores). Consequently, the generative process produces a CNA profile that aligns with the stronger class-level signals present in the conditioning modalities (RNA-Seq, WSI, RPPA). The resulting imbalance in classification accuracy should be interpreted as a marker of the generative models prioritizing global biological structures over the reconstruction of unique CNA information. The stage prediction task (Fig 2, Right) proved more challenging for all modalities, with smaller overall performance differences observed between the real and synthetic datasets.

This evaluation serves as a functional measure of generation quality. The outcomes confirm that our framework preserves existing predictive signals and that the generated data is not a superficial imitation but a meaningful synthesis of the original multimodal data.

### Data Generation to Mitigate Predictive Degradation in Multimodal Downstream Tasks

To demonstrate the downstream utility of our generative frameworks, we simulated a practical clinical challenge: applying a trained multimodal predictive model to new patients for whom some data modalities are unavailable. We first trained two separate models on the complete multimodal training set: a Random Forest classifier for tumor stage prediction and a Random Survival Forest [48] for survival analysis.

We then evaluated these trained models on the test set under numerous missing data conditions, where one or more modalities were removed. To isolate the predictive power beyond the primary cancer type signature, that previous results established as a dominant biological signal that is well-preserved in the generated data, we also included a baseline model trained only on the cancer type labels. This baseline demonstrates the performance attributable solely to knowing the average stage or survival outcome for a given cancer type, allowing us to measure the additional predictive power derived from the more refined, intra-tumor biological signals that our framework generates. We compared three outcomes for each condition: Ablation, representing the performance on the dataset with the modality (or modalities) missing; Synthetic from Multi-condition and Synthetic from Coherent Denoising, for our two reconstruction methods to generate missing data.

The results for the tumor stage classification (F1 Score) and survival analysis (C-index) tasks are presented in Fig 3, top and bottom panels respectively, while Table 2 provides a detailed quantitative breakdown of the performance changes. Detailed performance metrics, including balanced accuracy, are available in S4 Appendix.

**Table 2.** Quantitative Impact of Generative Completion on Downstream Task Performance. The table quantifies the change in performance for tumor stage classification (F1 Score) and survival analysis (C-Index) under various data sparsity scenarios. All values are reported as mean (±standard deviation) across 10 experimental runs. *“Drop from Ablation”* indicates the performance decrease when modalities are removed, relative to the baseline model that uses the complete dataset. *“Gain”* indicates the subsequent performance increase after synthetically generating the missing data, relative to the performance on the ablated (incomplete) data.

| Test Condition | Stage Classification (F1 Score) |  |  | Survival Analysis (C-Index) |  |  |
| --- | --- | --- | --- | --- | --- | --- |
|  | Ablation Drop | Gain from generated data |  | Ablation Drop | Gain from generated data |  |
|  |  | Multi-condition | Coherent Denoising |  | Multi-condition | Coherent Denoising |
| No Cna | -0.010 ± 0.012 | +0.008 ± 0.013 | +0.011 ± 0.009 | -0.025 ± 0.007 | +0.026 ± 0.007 | +0.028 ± 0.006 |
| No Rnaseq | -0.045 ± 0.012 | +0.044 ± 0.008 | +0.037 ± 0.012 | -0.163 ± 0.016 | +0.162 ± 0.016 | +0.160 ± 0.016 |
| No Rppa | -0.037 ± 0.007 | +0.036 ± 0.012 | +0.033 ± 0.012 | -0.021 ± 0.004 | +0.020 ± 0.005 | +0.022 ± 0.005 |
| No Wsi | -0.027 ± 0.008 | +0.021 ± 0.012 | +0.020 ± 0.009 | -0.047 ± 0.003 | +0.034 ± 0.003 | +0.037 ± 0.003 |
| No Cna, Rnaseq | -0.065 ± 0.020 | +0.058 ± 0.016 | +0.059 ± 0.022 | -0.187 ± 0.016 | +0.184 ± 0.018 | +0.186 ± 0.018 |
| No Cna, Rppa | -0.066 ± 0.014 | +0.058 ± 0.014 | +0.066 ± 0.015 | -0.035 ± 0.005 | +0.037 ± 0.007 | +0.040 ± 0.006 |
| No Cna, Wsi | -0.034 ± 0.009 | +0.031 ± 0.011 | +0.026 ± 0.011 | -0.082 ± 0.010 | +0.074 ± 0.009 | +0.075 ± 0.009 |
| No Rnaseq, Rppa | -0.098 ± 0.022 | +0.084 ± 0.021 | +0.083 ± 0.019 | -0.114 ± 0.011 | +0.107 ± 0.012 | +0.109 ± 0.011 |
| No Rnaseq, Wsi | -0.206 ± 0.025 | +0.187 ± 0.026 | +0.193 ± 0.027 | -0.197 ± 0.015 | +0.179 ± 0.015 | +0.185 ± 0.017 |
| No Rppa, Wsi | -0.066 ± 0.010 | +0.057 ± 0.013 | +0.055 ± 0.011 | -0.084 ± 0.005 | +0.072 ± 0.006 | +0.076 ± 0.007 |
| No Cna, Rnaseq, Rppa | -0.202 ± 0.024 | +0.176 ± 0.017 | +0.188 ± 0.021 | -0.117 ± 0.020 | +0.095 ± 0.021 | +0.107 ± 0.021 |
| No Cna, Rnaseq, Wsi | -0.149 ± 0.035 | +0.117 ± 0.036 | +0.136 ± 0.036 | -0.219 ± 0.016 | +0.190 ± 0.015 | +0.207 ± 0.017 |
| No Cna, Rppa, Wsi | -0.079 ± 0.020 | +0.069 ± 0.021 | +0.067 ± 0.019 | -0.100 ± 0.020 | +0.094 ± 0.018 | +0.097 ± 0.017 |
| No Rnaseq, Rppa, Wsi | -0.224 ± 0.013 | +0.114 ± 0.018 | +0.123 ± 0.021 | -0.173 ± 0.020 | +0.089 ± 0.019 | +0.101 ± 0.022 |

**Fig 3.**
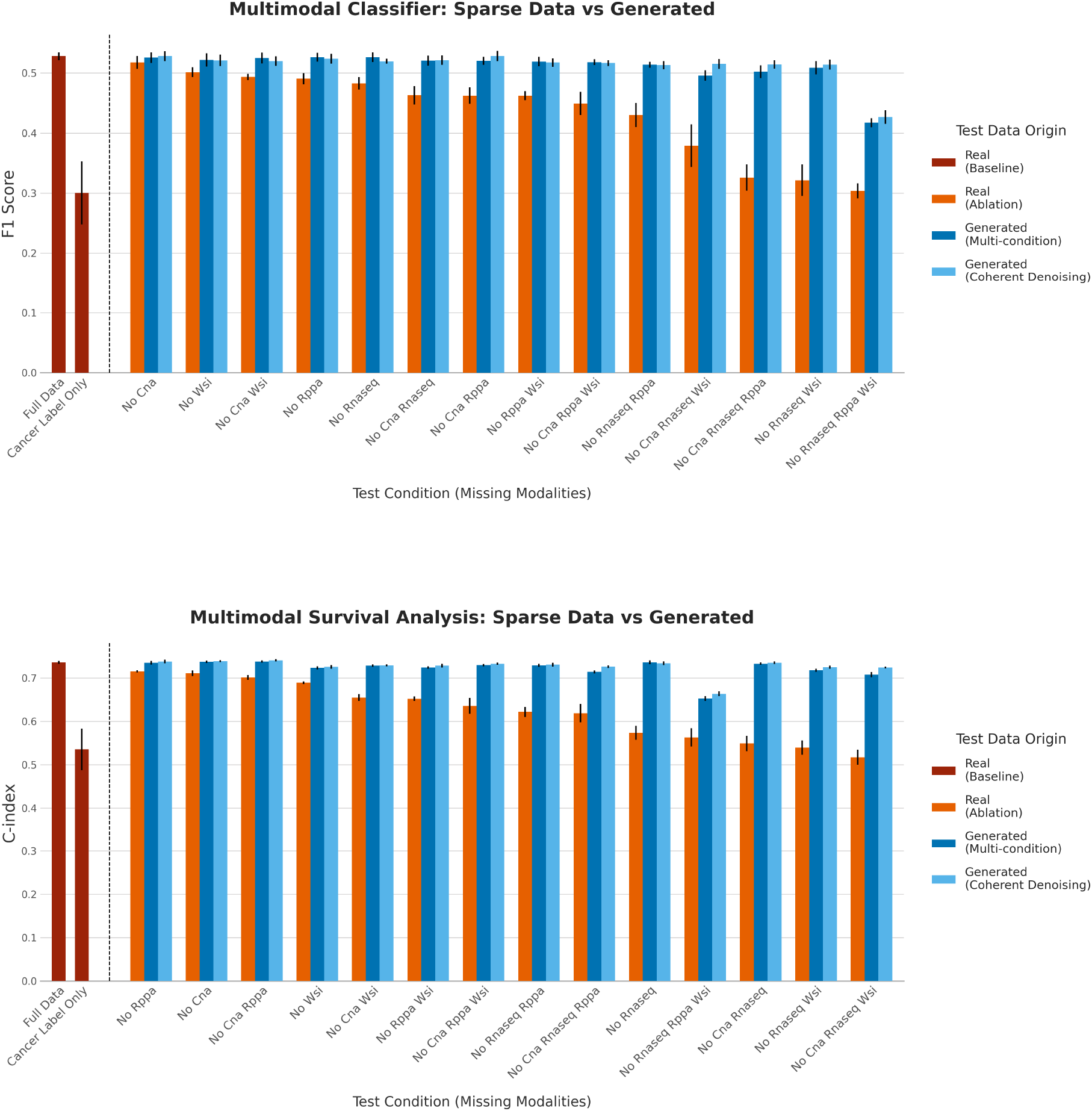
Performance comparison for multimodal downstream tasks on sparse versus synthetically generated data. The plots show results for a tumor stage classifier (top) and a survival model (bottom). We compare the performance on the complete test set (Baseline, dark red) against scenarios where modalities are removed (Ablation, orange) and where missing data is imputed by either the Synthetic Multi (dark blue) or Synthetic Coherent (light blue) models. As a secondary baseline, performances of the predictive model when only the cancer label is given as input are shown. Error bars represent the standard deviation across 10 experimental runs.

Across both tasks, the removal of modalities often leads to a significant degradation in predictive performance. As shown in Fig 3, the performance under these Ablation conditions can become substantially lower than the Full Data baseline. As quantified in Table 2, this drop is particularly severe when the most informative modalities are missing. For instance, removing both RNA-Seq and WSI data resulted in an F1-score decrease of 0.206 (± 0.025) for stage classification. For survival analysis, the C-Index dropped by as much as 0.219 (± 0.016) when CNA, RNA-Seq, and WSI data were all absent.

Crucially, in every tested sparse data scenario, generating the missing data using either the multi-condition or ensemble generation via Coherent Denoising models resulted in a marked recovery of performance. Post-hoc analysis confirmed that this performance improvement over the ablated baseline was statistically significant in all conditions for both stage classification and survival analysis (see full details in S6 Appendix, Section 2). The only exception is the scenario where only CNA data was missing for stage classification, where the ablated data already performs at the same level of the full dataset.

This performance rescue was particularly pronounced in cases of extreme data sparsity. For instance, in the survival analysis task (Fig 3, bottom panel), removing RNA-Seq, RPPA, and WSI data caused the C-index to collapse from a baseline of 0.736 to 0.563. Generating this missing data restored the C-index to approximately 0.66, recovering a substantial portion of the lost performance. Similarly, for stage classification (Fig 3, top panel), generating missing data consistently brought the F1 score much closer to the full data baseline than the ablated version. Notably, even when more than a single modality was missing, the performance with synthetically completed data was often statistically indistinguishable from that of the Full Data baseline (full details in S6 Appendix, Section 2), indicating a near-complete performance recovery.

A comparison between the Coherent Denoising and multi-condition models shows broadly comparable performance, with performance differences that were generally not statistically significant, suggesting neither framework was universally superior across all conditions. Both proved to be effective at generating high-utility data. In summary, these experiments demonstrate a key application of our generative framework: as an inference-time tool to complete sparse patient profiles, thereby mitigating the performance loss of downstream predictive models and enabling more robust analyses in the face of incomplete data.

### Counterfactual Analysis for Diagnostic Prioritization

As a final demonstration of utility, we designed an experiment to test whether our generative models could guide the efficient prioritization of diagnostic resources. We simulated a clinical scenario where a costly but informative modality (e.g., RNA-Seq or WSI) is not readily available and would need to be acquired. The objective is to determine if our models can help prioritizing patients that would benefit most from observing that additional modality in order to maximize the performance of a downstream predictive task, such as tumor stage classification.

To investigate this, we first trained a multimodal Random Forest classifier for tumor stage prediction. We then defined a counterfactual *variance score* for each patient in the test set. This score quantifies how much the classifier’s prediction changes when the missing real data for that modality is substituted with multiple different versions synthesized by our generative models (conditioned on the patient’s other available modalities). A low variance score indicates that the generated data consistently leads to the same prediction, suggesting the modality information for that patient is largely reconstructible from their other original data. A high score suggests the real data contains unique, non-redundant information that the model cannot infer.

We then compared two strategies for progressively acquiring the RNA-Seq or WSI modality for the test set, as shown in Fig 4: Random Prioritization, a baseline strategy where the data is observed for a randomly selected subset of patients; and Informed Prioritization, a strategy guided by our counterfactual variance score, where RNA-Seq and WSI data is preferentially observed for the patients with the highest variance scores first.

**Fig 4.**
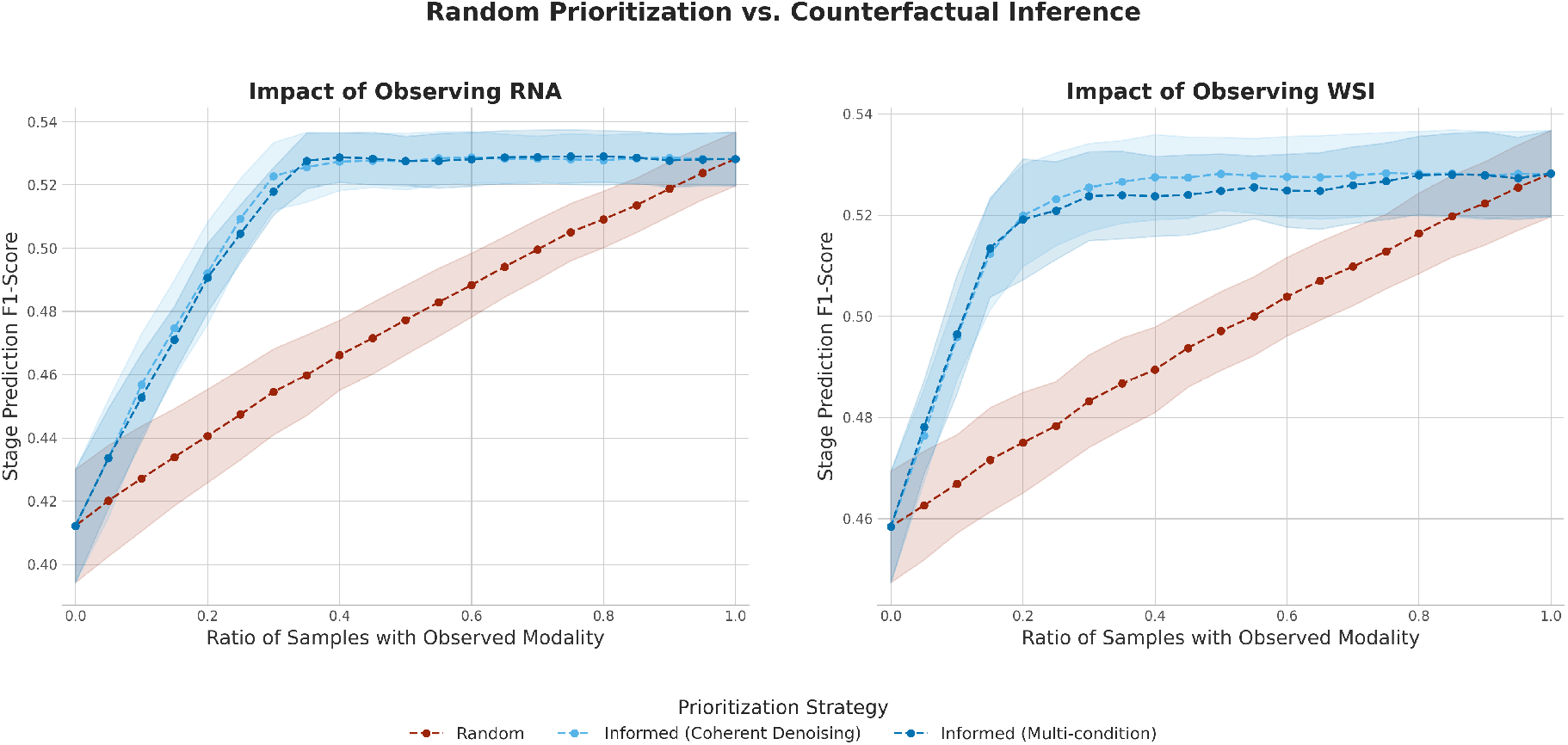
Evaluating Counterfactual Inference for Prioritizing RNA-Seq (Left) and WSI (Right) Data Acquisition. The plot shows the F1-score of a multimodal stage prediction classifier as the ratio of patients with an observed modality (RNA-Seq or WSI data) is varied. The Random Prioritization strategy (red) removes that modality data from patients at random. The Informed Prioritization strategies (blue) use a counterfactual variance score to preferentially acquire that modality data for the most informative patients first. Error bands show the standard deviation across 10 experimental repetitions. Note that the two plots are on separate y-axis scales, because of the intrinsic difference in performance that a classifier has with and without that modality.

The results demonstrate the clear benefit of the counterfactual-guided approach. The Random Prioritization strategy (Fig 4, red line) serves as a baseline, showing a near-linear increase in classifier performance (F1-score) as the proportion of samples with the observed modality increases from 0% to 100%. In contrast, the Informed Prioritization strategy (blue lines) yields a much more rapid performance gain, reaching a near-optimal F1-score even when the modality is acquired for only a fraction of the cohort. For instance, when observing RNA-Seq data (left panel), the informed strategy achieves near-peak performance with only 40% of samples observed—a level of performance that the random strategy only reaches after observing more than 90% of the cohort. The overall superiority of both informed prioritization strategies compared to the random baseline was statistically significant (see full details in S6 Appendix, Section 3). We validated this finding for the tumor stage classification task (Fig 4) and additionally confirmed its robustness by extending the analysis to survival prediction, where the informed strategy similarly outperformed the random baseline (see S5 Appendix). This demonstrates that the counterfactual variance score effectively identifies the small subset of patients for whom the target modality is most critical, allowing for a highly efficient data acquisition strategy.

This experiment serves as a proof-of-concept for a powerful application of cross-modal generative models. By identifying patients for whom a given modality is most impactful, these models can provide a quantitative framework to navigate the complex landscape of modern diagnostics. In a clinical reality where hundreds of available tests can lead to patient wait times of weeks or months [49], such a framework allows for the strategic prioritization of the most critical assays. This, in turn, maximizes the utility of finite resources, shortens the path to a definitive diagnosis, and ensures that clinical decisions are made in a more timely and informed manner.

### Privacy Preservation Properties in Unconditional Generation

A critical consideration for generative models trained on sensitive patient data is the potential for training data reconstruction. To assess this risk, we evaluated the behavior of our two frameworks under an unconditional generation setting, where no conditioning modalities were provided as input at inference time. For the multi-condition model, this involved providing a fully masked (null) condition. For the Coherent Denoising ensemble, the single-condition models (which were never trained for unconditional generation) were fed a null zero vector as their condition. This experiment tests whether the models have memorized the training distribution to a degree that would allow the recreation of patient-like profiles without specific input, posing a privacy concern. In this context, a failure to generate a variable, realistic data manifold is the desired, privacy-preserving outcome.

The results are visualized through UMAP and PCA projections in Fig 5 and quantified in Table 3 through F1-score [50] - for coverage - and Energy Distance (ED) [51]. They reveal a significant difference between the two approaches. The multi-condition model, which is trained using a masking strategy that includes empty conditioning sets, learns to reconstruct a substantial portion of the training data manifold even without explicit inputs (Fig 5, left panels). This is confirmed quantitatively by its low Energy Distance of 1.01 and a non-null F1 score of 0.14, indicating it has learned a strong internal representation of the overall data distribution.

**Table 3.** Quantitative Assessment of Unconditional Generation and Privacy. Comparison of the two models on their ability to reconstruct the training data manifold without conditioning. We report the *F1 Score* (higher indicates better manifold reconstruction, worse for privacy) and the *Energy Distance* (lower is more similar, worse for privacy) between the generated and real distributions. Values are mean ± std over 10 runs.

| Model | F1 Score | Energy Distance |
| --- | --- | --- |
| Multi-condition | 0.1423 ± 0.0117 | 1.0076 ± 0.0083 |
| Coherent Denoising | 0.0000 ± 0.0000 | 2.1056 ± 0.0062 |

**Fig 5.**
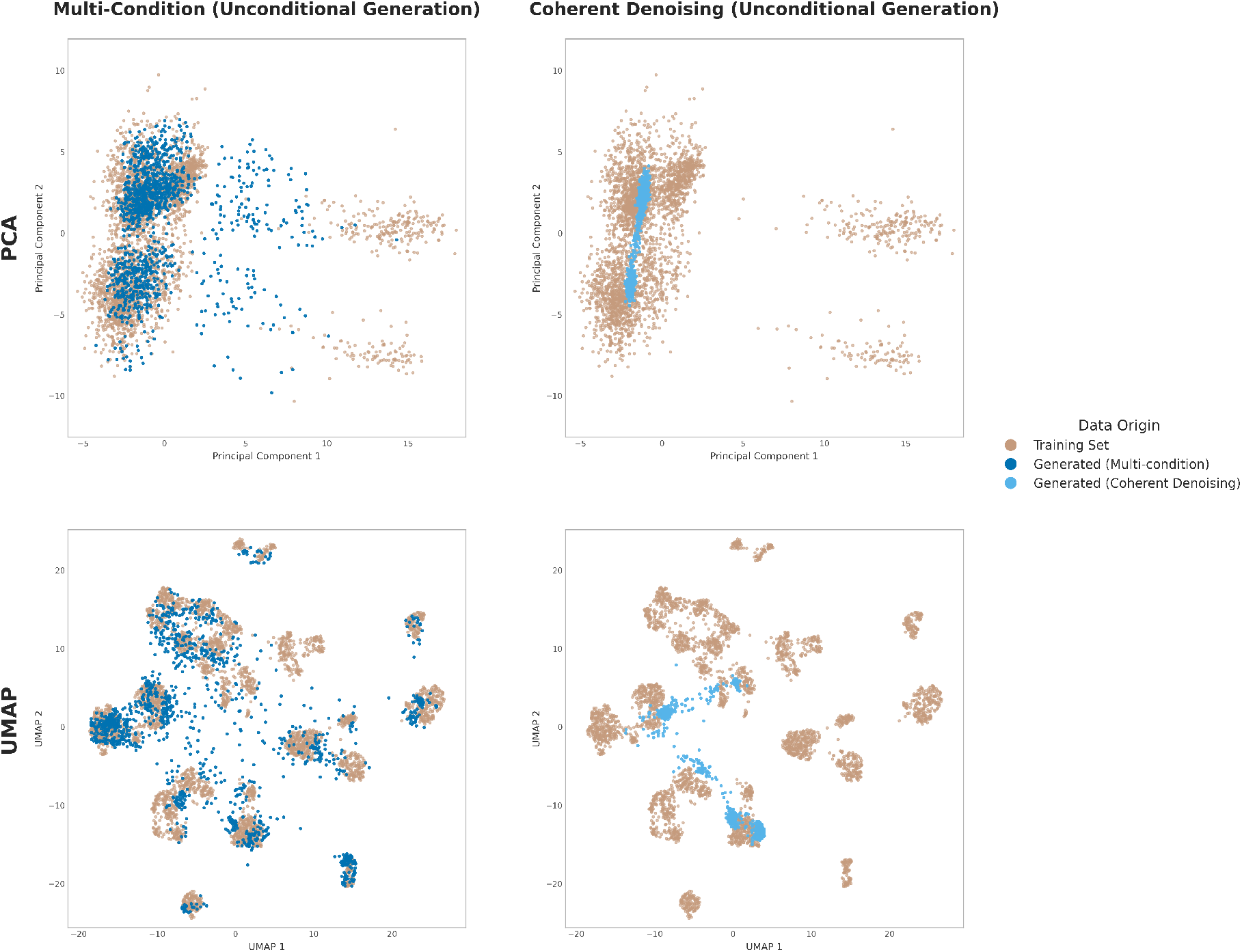
Privacy Preservation in Unconditional Generation. Qualitative comparison of the training set manifold with data generated unconditionally by the multi-condition (left panels) and Coherent Denoising (right panels) models. PCA (top) and UMAP (bottom) projections show the ability of the multi-condition model (that is trained also on masked data) to learn and reconstruct a good part of the data distribution even without any specific conditioning at inference time. On the other hand our Coherent Denoising approach is only able to produce unrealistic samples around the mean of the distribution. This is a highly desirable property in contexts where privacy preservation of the training set is crucial.

In contrast, the Coherent Denoising ensemble is inherently robust against such reconstruction. Since it is composed of models trained only on single-condition pairs, it cannot generate realistic data without conditioning if this approach has not been explicitly enabled during training. In the unconditional setting, the ensemble produces unrealistic samples tightly clustered around the distribution’s mean, failing to replicate the specific, separable clusters of the training data (Fig 5, right panels). This is evidenced by a high Energy Distance of 2.11 and an F1 score of zero, showing its output is structurally dissimilar to the training data and fails to recapitulate the training set manifold.

This outcome demonstrates a key privacy-preserving advantage of the Coherent Denoising framework. Its capability to explicitly enforce conditioning for meaningful generation enables additional safety layers and inherently mitigates the risk of inadvertently exposing sensitive training data, making it a more secure choice for applications in clinical settings where data privacy is crucial.

## Discussion

The integration of multimodal data is a central goal across biomedicine, as models that fuse multiple data types consistently outperform single-modality approaches for understanding complex biological systems [7, 8]. A common and effective strategy for this integration, adopted in this work, involves encoding heterogeneous data into a unified, low-dimensional embedding space. This approach has been significantly advanced by foundation models capable of learning powerful data representations [9, 10, 12]. However, the clinical translation of such integrative models is limited by the practical challenge of data sparsity. Patient datasets are frequently incomplete due to the cost, availability, and logistical complexity of data acquisition [15, 16, 17, 18, 19]. While generative AI, particularly state-of-the-art Diffusion Models, has shown success in single-modality and narrowly-defined conditional synthesis tasks [36, 39, 40], current approaches remain too confined to address the challenge of general, any-to-any multimodal data generation for biomedical research.

In this study, we developed a generative AI framework to address data sparsity in biomedical datasets. A key feature of our approach is its comprehensive multimodal capability: it is designed not to generate a single target but to synthesize any of four major data types - CNA, RNA-Seq, RPPA, and WSI embeddings. To achieve this flexibility, we implemented and benchmarked two distinct diffusion-based strategies. The first, a multi-condition model, represents a monolithic approach where a single, large network learns to condition on an arbitrary set of inputs via a masking mechanism. As an advantageous alternative, we introduce ensemble generation via Coherent Denoising, a novel method that offers significant benefits in modularity and scalability. This technique operates as a flexible ensemble of simpler, independently trained, single-condition models. The modular design is inherently more scalable. For example, incorporating a new data modality, only requires training new pairwise models without altering the existing validated components of the framework. The effectiveness of both architectures was demonstrated on a large-scale, aggregated dataset of over 10,000 multimodal samples from the TCGA program across 20 different cancer types.

We performed a multi-step validation of generation quality. This included qualitative confirmation of data manifold preservation via UMAP, quantitative measurement of reconstruction fidelity, and functional tests of predictive signal preservation in single-modality classifiers. Our key findings highlight the downstream utility of the synthetic data produced by our framework. We’ve shown that this data can be leveraged during inference to reconstruct sparse patient profiles, thereby enhancing the performance of multimodal models for critical cancer research applications, specifically tumor staging and survival analysis.

Across all tested sparse data conditions, our generative completion yielded a significant performance improvement over using the incomplete data alone, often achieving results with no significant difference to that of the model using the complete, original dataset. This capability relies on the preservation of key biological signals that extend beyond the dominant tumor-type signature, as our full data models consistently outperformed baselines that used only cancer type information. Furthermore, we established a novel application of these models in counterfactual inference, providing a quantitative method to guide the prioritization of diagnostic data acquisition by identifying patients for whom a specific modality would be most informative.

Beyond the specific application in this study in a controlled multi-omic, multimodal setting, the ensemble generation via Coherent Denoising framework is intended to integrate insights from a much broader, heterogeneous ecosystem of models. This offers a practical solution to the significant challenge of creating a single, all-encompassing model to handle every type of patient data, from genomics and imaging to phenotype and exposome.

The Coherent Denoising framework is designed to combine information from many different and specialized models, not just the specific data used in this study. This approach is particularly well-suited for healthcare, where the diversity of data modalities favors a collection of domain-specific models over a single, all-encompassing one. Our framework provides a principled method for these specialized expert models to contribute to a unified generative task by forming a consensus. This makes it a prime candidate for synthesizing a truly holistic patient view by leveraging a ‘community of experts’ rather than attempting to build one all-knowing model. The modular design also confers significant privacy advantages, as the ensemble is inherently robust against unintentional data reconstruction, a critical feature for models handling sensitive clinical information.

This work points toward a long-term vision for data-driven approaches in both biomedical research and clinical practice. The ability to conditionally generate high-fidelity data provides a conceptual step towards future in silico experimentation, where the entire patient’s manifold could be represented by sampling multiple multimodal profiles for the purpose of testing novel biomarkers or simulating patient states under hypothetical perturbations. In the clinical setting, our counterfactual inference analysis showcases a preliminary application of personalized medicine through adaptive diagnostic workflows, where clinical testing strategies can be tailored to an individual patient to maximize information gain.

The models were trained and validated exclusively on data from the TCGA project; external validation on independent clinical cohorts will be critical to ensure generalizability. We demonstrated the potential of our framework on four selected modalities, but this could be expanded to additional data types, such as DNA methylation or metabolomics, which were not available or too limited in the considered dataset. We selected a highly compressed latent dimension to demonstrate that the framework is robust in a challenging, information-dense scenario. Future work will explore the impact of varying embedding dimensions, which might retain a greater biological signal. Lastly, the downstream utility assessment could also be expanded to other tasks such as treatment response.

## Methods

### Data and Code Availability

The TCGA data used in this work was downloaded from the GDC Data Portal [52] and the UCSC Xena Hub [53]. The code used to perform the data preprocessing, train the generative models, and reproduce the results in this paper is available on GitHub at: https://github.com/r-marchesi/coherent-genAI. All experiments were conducted on FBK’s high-performance computing cluster (Abacus), utilizing NVIDIA H100 (80GB) and L40s (48GB) GPUs. Nodes were equipped with Intel Xeon Platinum 8470 CPUs and had access to at least 200GB of system RAM. Training for each individual model (20,000 epochs) typically concluded within 24-hours. The main pipeline of our methodological framework is presented in Fig 6.

**Fig 6.**
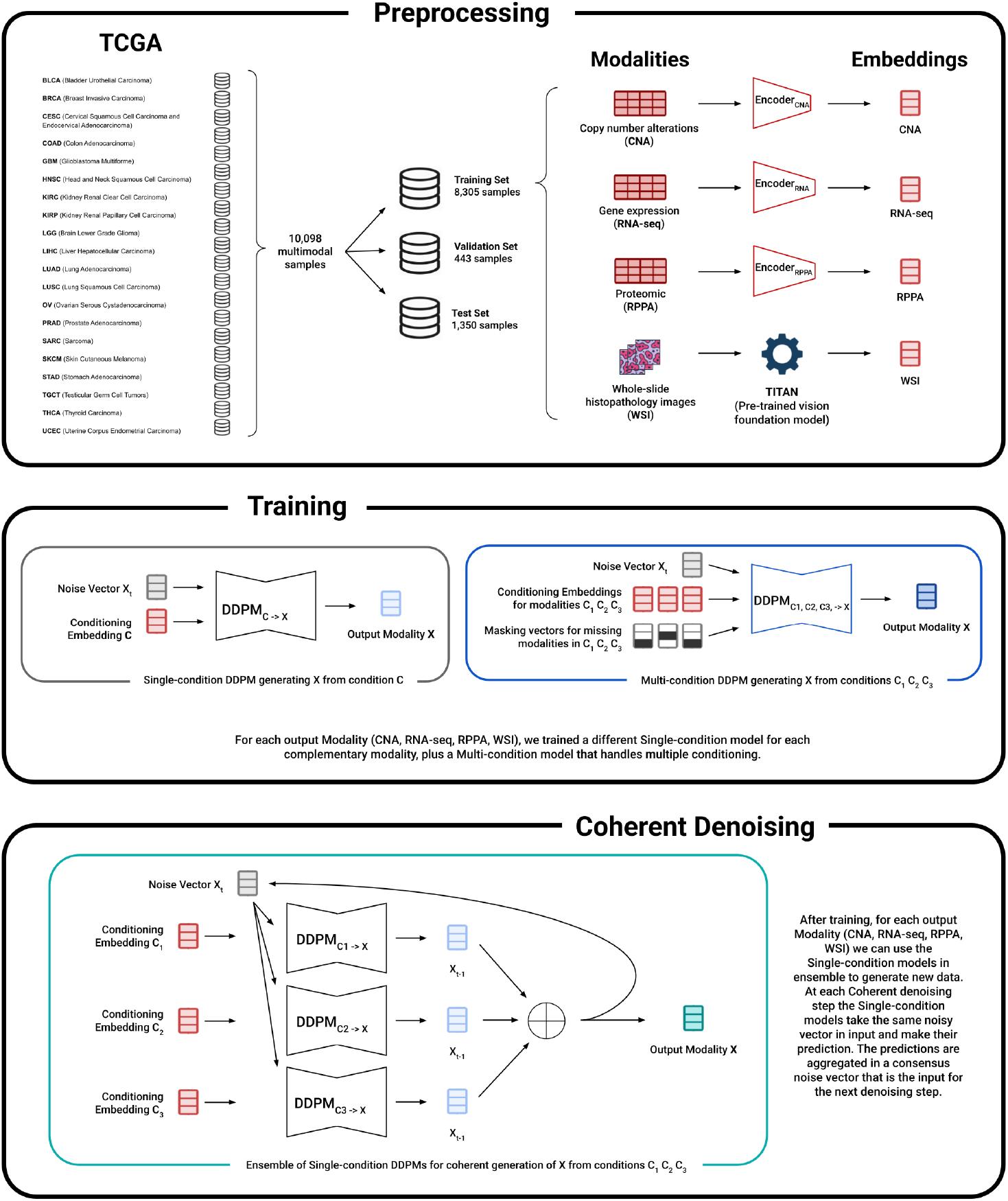
Overview of the Methodological Pipeline. (Top Panel: Preprocessing) Data from the TCGA cohort, spanning 20 cancer types, is curated for the study. The omics modalities (CNA, RNA-seq, RPPA) are encoded into a harmonized 32-dimensional latent space using modality-specific trained autoencoders, while Whole-Slide Image (WSI) data is first embedded using the Titan foundation model and then reduced to the same dimension via Principal Component Analysis (PCA). (Middle Panel: Training) Two types of DDPMs are trained on these embeddings: single-condition models for one-to-one generation (*C*→ *X*), and a masked multi-condition model for many-to-one generation (*C*_1_,*C*_2_,*C*_3_ →*X*). (Bottom Panel: Coherent Denoising) Our novel ensemble strategy utilizes the collection of pre-trained single-condition models to generate a target modality at inference time. During the reverse diffusion process, at each denoising step *t*, every model makes an independent prediction based on the same input vector *X*_*t*_. These individual noise predictions are then aggregated, into a single consensus vector. This consensus noise then guides the subsequent denoising step, effectively forcing the generation to satisfy multiple conditions coherently throughout the entire process. The icons in the top panel were created using generative artificial intelligence (Google Gemini) and are used in accordance with Google’s Terms of Service (https://policies.google.com/terms).

### Data Collection and Preprocessing

Tumor multi-omic data were retrieved from The Cancer Genome Atlas (TCGA) via the UCSC Xena platform, comprising a total of 10,098 primary tumor samples across 20 distinct cancer types. Details about dataset composition are presented in Table 4.

**Table 4.** Data Summary. Each row corresponds to a cancer type included in the study. The table reports the total number of tumor samples, and for each modality (CNA, RNA-seq, Proteomics, WSI), the number and percentage of samples available. “Complete” indicates samples with all four modalities available; “Incomplete” includes all samples with one or more missing modalities.

| Cancer Type | Total Samples | CNA | RNA | Prot | WSI | Complete | Incomplete |
| --- | --- | --- | --- | --- | --- | --- | --- |
| BRCA | 1224 | 1057 (86.4%) | 1214 (99.2%) | 919 (75.1%) | 1054 (86.1%) | 812 (66.3%) | 412 (33.7%) |
| KIRC | 610 | 521 (85.4%) | 606 (99.3%) | 477 (78.2%) | 513 (84.1%) | 442 (72.5%) | 168 (27.5%) |
| UCEC | 584 | 537 (92.0%) | 581 (99.5%) | 440 (75.3%) | 504 (86.3%) | 393 (67.3%) | 191 (32.7%) |
| LUAD | 581 | 507 (87.3%) | 577 (99.3%) | 365 (62.8%) | 468 (80.6%) | 310 (53.4%) | 271 (46.6%) |
| HNSC | 574 | 517 (90.1%) | 566 (98.6%) | 354 (61.7%) | 450 (78.4%) | 315 (54.9%) | 259 (45.1%) |
| LUSC | 555 | 497 (89.5%) | 551 (99.3%) | 328 (59.1%) | 477 (85.9%) | 301 (54.2%) | 254 (45.8%) |
| PRAD | 551 | 474 (86.0%) | 550 (99.8%) | 352 (63.9%) | 349 (63.3%) | 231 (41.9%) | 320 (58.1%) |
| OV | 530 | 494 (93.2%) | 380 (71.7%) | 385 (72.6%) | 101 (19.1%) | 50 (9.4%) | 480 (90.6%) |
| LGG | 530 | 520 (98.1%) | 530 (100.0%) | 435 (82.1%) | 493 (93.0%) | 400 (75.5%) | 130 (24.5%) |
| GBM | 523 | 498 (95.2%) | 154 (29.4%) | 218 (41.7%) | 338 (64.6%) | 42 (8.0%) | 481 (92.0%) |
| COAD | 503 | 447 (88.9%) | 501 (99.6%) | 363 (72.2%) | 432 (85.9%) | 331 (65.8%) | 172 (34.2%) |
| STAD | 478 | 429 (89.7%) | 448 (93.7%) | 357 (74.7%) | 347 (72.6%) | 255 (53.3%) | 223 (46.7%) |
| SKCM | 475 | 463 (97.5%) | 473 (99.6%) | 352 (74.1%) | 211 (44.4%) | 159 (33.5%) | 316 (66.5%) |
| THCA | 460 | 0 (0.0%) | 458 (99.6%) | 304 (66.1%) | 407 (88.5%) | 0 (0.0%) | 460 (100.0%) |
| BLCA | 432 | 403 (93.3%) | 425 (98.4%) | 343 (79.4%) | 299 (69.2%) | 244 (56.5%) | 188 (43.5%) |
| LIHC | 429 | 365 (85.1%) | 423 (98.6%) | 184 (42.9%) | 339 (79.0%) | 157 (36.6%) | 272 (63.4%) |
| KIRP | 324 | 284 (87.7%) | 323 (99.7%) | 216 (66.7%) | 262 (80.9%) | 190 (58.6%) | 134 (41.4%) |
| CESC | 312 | 296 (94.9%) | 309 (99.0%) | 172 (55.1%) | 196 (62.8%) | 121 (38.8%) | 191 (61.2%) |
| SARC | 267 | 247 (92.5%) | 264 (98.9%) | 226 (84.6%) | 224 (83.9%) | 181 (67.8%) | 86 (32.2%) |
| TGCT | 156 | 155 (99.4%) | 156 (100.0%) | 122 (78.2%) | 154 (98.7%) | 120 (76.9%) | 36 (23.1%) |
| **TOTAL** | **10098** | **8711 (86.3%)** | **9489 (94.0%)** | **6912 (68.4%)** | **7618 (75.4%)** | **5054 (50.0%)** | **5044 (50.0%)** |

The study integrated five data modalities. Clinical/diagnostic information was used exclusively for downstream validation. Gene expression data were obtained from RNA sequencing (RNA-seq). Proteomic data were derived from Reverse Phase Protein Array (RPPA) assays. Copy number alterations (CNA) were provided at the gene level. Histological features were extracted from whole-slide histopathology images (WSI) using Titan [11], a pre-trained vision foundation model that generates fixed-length slide-level embeddings from raw WSI. Titan uses Conch 1.5 [54] as patch encoder, which crops patches of 512×512 pixels at 20 × magnification. Titan outputs a slide embedding of 736×1 dimension. Inference was performed using the Trident [55, 56] package.

### Preprocessing

Gene expression data were initially provided as log_2_(count + 1) values. These were exponentiated to recover raw count estimates, normalized to Counts Per Million (CPM) to adjust for library size variation, and log-transformed again as log_2_(CPM+1) to stabilize variance. Genes were retained if CPM *>* 1.0 in at least 20% of samples, in line with standard filtering practices to exclude low-abundance transcripts. Gene-level copy number values were derived from ABSOLUTE-based segmentations [57]. Values equal to 0 were excluded, as they are undefined under log_2_ transformation and can bias downstream analyses. Remaining values were transformed to log_2_ ratios relative to the diploid state (log_2_[CNA / 2]) and clipped to the range [–2, 2] to reduce the influence of extreme events and outliers. RPPA values were median-centered by subtracting, for each protein, the median expression across all samples to remove sample-independent offsets and center each distribution. All modalities underwent standardized quality control: biological outliers were removed based on modality-specific distributions, and features with more than 10% missing values were excluded. For the remaining data, RNA-seq and RPPA missing values were imputed using K-nearest neighbors (KNN), while CNA missing values were imputed using the feature-wise median.

### Data Splitting and Scaling

The dataset was stratified by cancer type and split into training (80%, 8,305 samples), validation (5%, 443 samples), and test (15%, 1,350 samples). This large, held-out test set provides a robust and computationally feasible evaluation for deep learning frameworks of this scale, serving as a standard alternative to k-fold cross-validation which was deemed computationally prohibitive for this study. Only samples with all modalities available were included in the test set in order to provide a complete and robust ground-truth for evaluating reconstruction fidelity and downstream task performance across modalities. All normalization, scaling and imputation procedures were applied after the split to prevent data leakage.

### Autoencoder embedding

To create a dense and low-dimensional representation, a separate autoencoder was trained for each modality (CNA, RNA-Seq, RPPA) on the training set. Each autoencoder learned to compress its respective modality into a 32-dimensional latent representation, a dimension selected to balance information density with model complexity. For WSI data, which were already embedded, Principal Component Analysis (PCA) was used to reduce their dimensionality.

To validate the robustness of our framework beyond the selected 32-dimensional latent space, we conducted an analysis across different embedding dimensions. As detailed in S10 Appendix, increasing dimensionality did not produce performance gains for the real data on downstream tasks, confirming that 32 dimensions sufficiently capture the available biological signal for this cohort size. Crucially, the generative framework maintained robust reconstruction fidelity and downstream utility even in higher-dimensional spaces, demonstrating its scalability to larger embedding models.

### Generative Diffusion Models

Our generative framework to synthesis of diverse biological modalities is built upon Denoising Diffusion Probabilistic Models (DDPMs) [32, 33], which learn to reverse a fixed process that gradually adds Gaussian noise to data.

Diffusion models operate by iteratively corrupting data with Gaussian noise and then learning to reverse this process to generate new samples. Formally, given a data sample *x*_0_ and a total of *T* noising steps, in the forward (noising) process, a fixed variance schedule 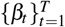 with *β*_*t*_ ∈ (0, 1) is defined. At each step *t*, Gaussian noise is added according to:

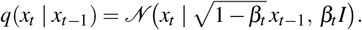

This process allows direct sampling of *x*_*t*_ from *x*_0_:

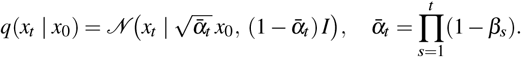

Here, *α*_*t*_ = 1 −*β*_*t*_ and 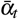 represents the cumulative signal retention up to step *t*.

The reverse (denoising) process is parameterized by a neural network *ε*_*θ*_ (*x*_*t*_, *t*) which predicts the noise component added at step *t*. The reverse transition probability is defined as:

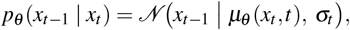

where the mean *µ*_*θ*_ (*x*_*t*_, *t*) is given by:

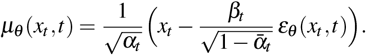

For practical implementations, we set *σ*_*t*_ = *β*_*t*_*I*.

Training involves minimizing a simplified variational bound, which effectively translates to matching the predicted noise with the true noise. The objective function is:

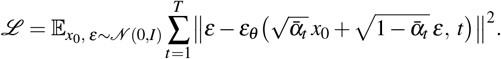

The term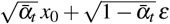 represents a sample from *q*(*x*_*t*_ *x*_0_), thereby enabling the network to learn the noise *ε* added at step *t*.

During inference, a pure noise sample *x*_*T*_ ∼ 𝒩 (0, *I*) is drawn, and the learned reverse transitions *p*_*θ*_ (*x*_*t*−1_ | *x*_*t*_) are applied iteratively for *t* = *T, T* − 1,…, 1 to yield a final sample *x*_0_ from the learned data distribution.

### Single-Condition Diffusion Models

To generate a target modality *X* conditioned on a single source modality *C*, we trained a separate diffusion model for each ordered pair of modalities. At each reverse step *t*, the neural network predicts the noise component *ε*_*θ*_ (*x*_*t*_, *t,C*). The models are optimized using a mean-squared error (MSE) objective:

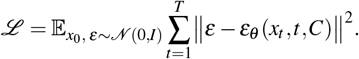

### Multi-Condition Diffusion Models

Extending the single-condition framework, a multi-condition diffusion model is capable of handling many conditioning modalities simultaneously. Each input modality *C*_*i*_ is first transformed via a linear projection. A masking strategy is implemented to enable the network to operate regardless of input availability: if a modality *C*_*i*_ is missing, its projected activation is set to zero, effectively nullifying its contribution to the subsequent layers of the network. The resulting projected vectors 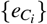 are concatenated and provided as input to the noise-prediction network 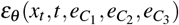. Training utilizes the same MSE objective as the single-condition models.

### Model Implementation and Training

We evaluated both multi-layer perceptrons (MLPs) and U-Net [58] as candidate architectures for all diffusion models. Contrary to their widespread use in image-based applications, U-Nets did not demonstrate superior performance over MLPs for our tabular embedding data. Consequently, all reported results were obtained using the simpler and more computationally efficient MLP architecture.

Our MLP-based noise prediction network processes three inputs: the noisy data (*x*_*t*_), the timestep (*t*), and one or more conditioning vectors (*C*_*i*_). Time is encoded using standard sinusoidal positional embeddings [59], and each conditioning vector is independently projected into a dedicated embedding space via a linear layer. The resulting embeddings are then concatenated with the noisy data (*x*_*t*_) to form the initial input to the network. Each hidden layer of the MLP consists of a linear transformation followed by Batch Normalization and a ReLU activation function. The sinusoidal time embedding is re-injected by concatenating it with the output of each hidden layer, ensuring the time signal is preserved throughout the network.

For hyperparameter optimization, we performed a comprehensive grid search. Key hyperparameters included learning rate ([10^−4^, 10^−3^]), batch size ([64, 128]), number of MLP layers ([4, 5, 6, 7]), MLP hidden size ([256, 512, 1024]), and dimensions for both time ([64, 128]) and conditional embeddings ([8, 16, 32]). Optimal parameters for each model were selected based on validation performance and are listed in S7 Appendix.

All models were trained for up to 20,000 epochs with early stopping. The stopping criterion was based on the Mean Squared Error (MSE) between fully denoised generated samples and their real counterparts on a held-out validation set. The model checkpoints with the best validation scores were saved for all subsequent analyses.

### Coherent Denoising

We introduce a novel ensemble generation technique termed **Coherent Denoising**. This approach contrasts with single, monolithic network architectures by instead leveraging a flexible ensemble of pre-trained, single-condition diffusion models. The ensemble generates a target modality *X* by aggregating evidence from multiple available conditioning modalities {*C*_1_,*C*_2_,…,*C*_*N*_}.

The method is integrated directly into the iterative reverse diffusion process. The generation begins by initializing a sample with pure Gaussian noise, *x*_*T*_ ∼𝒩 (0, **I**). Then, for each denoising step *t* from *T* down to 1, each model *M*_*i*_ in the ensemble predicts a noise component, *ε*_*θi*_ (*x*_*t*_,*C*_*i*_, *t*), based on the current noisy sample *x*_*t*_ and its corresponding conditioning data *C*_*i*_. These individual predictions are aggregated into a single consensus noise vector, *ε*_consensus_, via a weighted average:

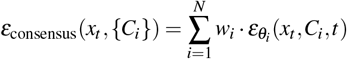

where the weights *w*_*i*_ are non-negative, sum to one, and can be set to reflect model reliability (e.g., inversely proportional to validation loss). For all reported experiments, we used a performance-based weighting scheme where weights were set inversely proportional to each single-condition model’s reconstruction MSE on the held-out validation set. This consensus noise is then used in the denoising update rule to compute the less-noisy sample *x*_*t*−1_. S8 Appendix presents a weight distribution analysis for all our models.

The underpinning of this ensemble approach is motivated by the connection between diffusion models and score-based generative modeling [60]. The noise prediction, *ε*_*θ*_ (*x*_*t*_,*C, t*), is trained to be proportional to the score of the data distribution, ∇_*xt*_ log *p*_*t*_(*x*_*t*_ |*C*). While a formal composition of scores using Bayes’ theorem would also require an unconditional score term [60], our method employs a practical heuristic that approximates the joint conditional score with a weighted average of the individual conditional scores. Therefore, the weighted average of noise predictions, *ε*_consensus_, serves as a computationally efficient proxy for the score of the joint distribution *p*_*t*_(*x*_*t*_ |*C*_1_,…,*C*_*N*_), avoiding the need to train and evaluate a separate unconditional model. This heuristic guides the reverse process toward a sample that simultaneously satisfies all conditions, with its efficacy demonstrated empirically in our results.

A key challenge in ensembling generative models is ensuring the constituent models do not provide conflicting guidance, which can degrade sample quality. We address this by monitoring the geometric agreement of the predicted noise vectors through a process of **coherence-based rejection sampling**. The vector 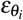 represents the denoising direction proposed by model *M*_*i*_; high angular alignment among these vectors indicates “coherence”. We quantify this using the pairwise cosine distance, 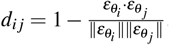. A generation trajectory is considered to have failed convergence and is rejected if the weighted average of these pairwise distances exceeds a predefined threshold (e.g., cosine distance = 1) for more than a small fraction (e.g., 5%) of the total denoising timesteps. This acts as a quality filter, discarding samples resulting from conflicting conditional evidence. We confirmed the stability of this threshold through a sensitivity analysis across a wide range of threshold values (±30%) in S9 Appendix, showing how the rejection rate is robust to minor changes in the threshold.

### Downstream Task Implementation

All downstream predictive models (Random Forest classifiers and Random Survival Forest) were trained using implementations that natively support missing values. For experiments in the “Data Generation to Mitigate Predictive Degradation” section, the Ablation condition involved passing input vectors to the trained models with missing modality features flagged as missing. For the Synthetic conditions, these missing features were first replaced with the generated data before being passed to the model. The Counterfactual Analysis experiment used a single multimodal classifier trained on all four modalities.

The counterfactual *variance score* for a patient (with a target modality, e.g., RNA-Seq, considered “missing”) was defined by a multi-step process: (1) Generate *N* = 10 different versions of the missing modality by running the generative model 10 times with different random seeds. (2) Pass these 10 complete, counterfactual patient profiles into the trained classifier, yielding 10 separate prediction probability vectors. (3) The score is the average variance of the class probabilities across these 10 predictions. A high score indicates prediction instability, implying the real data contains a predictive signal that is not redundant with the other available modalities. The plots in Fig 4 show the progression of the classifier’s F1-score as the real modality is progressively “acquired” for the test set population. The Random Prioritization (red line) shows the baseline performance as patients are randomly selected for acquisition. The Informed Prioritization (blue line) first sorts all patients by their variance score, from highest to lowest. It then shows the F1-score progression as the real modality is acquired in this informed order, demonstrating that acquiring the modality for the highest-variance patients first yields a much faster improvement in performance.

## Data Availability

The TCGA data used in this work was downloaded from the GDC Data Portal https://portal.gdc.cancer.gov/ and the UCSC Xena Hub https://xenabrowser.net/. The code used to perform the data preprocessing, train the generative models, and reproduce the results in this paper is available on GitHub at: https://github.com/r-marchesi/coherent-genAI.

## Author contributions statement

G.J. and V.O. conceived and supervised the study. N.L. conceived and developed the Coherent Denoising methodology. R.M. designed and implemented the main computational pipeline and models, with substantial support from W.E. on the codebase.

M.P. contributed to the implementation for WSI data handling. G.L. collected, curated, and preprocessed the datasets. R.M. and W.E. conducted the experiments and analyzed the results. F.R., S.B., and M.M. provided scientific guidance and oversight.

R.M. wrote the initial draft of the manuscript. R.M, F.R., W.E, S.B., and N.L. wrote the final version of the manuscript. All authors reviewed, edited, and approved the final manuscript.

## Supporting information

**S1 Appendix Distribution Fidelity for each Modality**

**S2 Appendix Full Reconstruction Accuracies**

**S3 Appendix Preservation of Predictive Signals in Generated Data**

**S4 Appendix Inference-Time Data Synthesis for Downstream Predictive Tasks**

**S5 Appendix Counterfactual Analysis for Survival Analysis**

**S6 Appendix Statistical Tests**

**S7 Appendix Hyperparameters**

**S8 Appendix Coherent Denoising Ensemble Weights**

**S9 Appendix Sensitivity Analysis of Rejection Sampling**

**S10 Appendix Impact of Embedding Dimension**

