## Supporting Information for "Coherent Cross-modal Generation of Synthetic Biomedical Data to Advance Multimodal Precision Medicine"

### S1 Appendix: Distribution Fidelity for each Modality

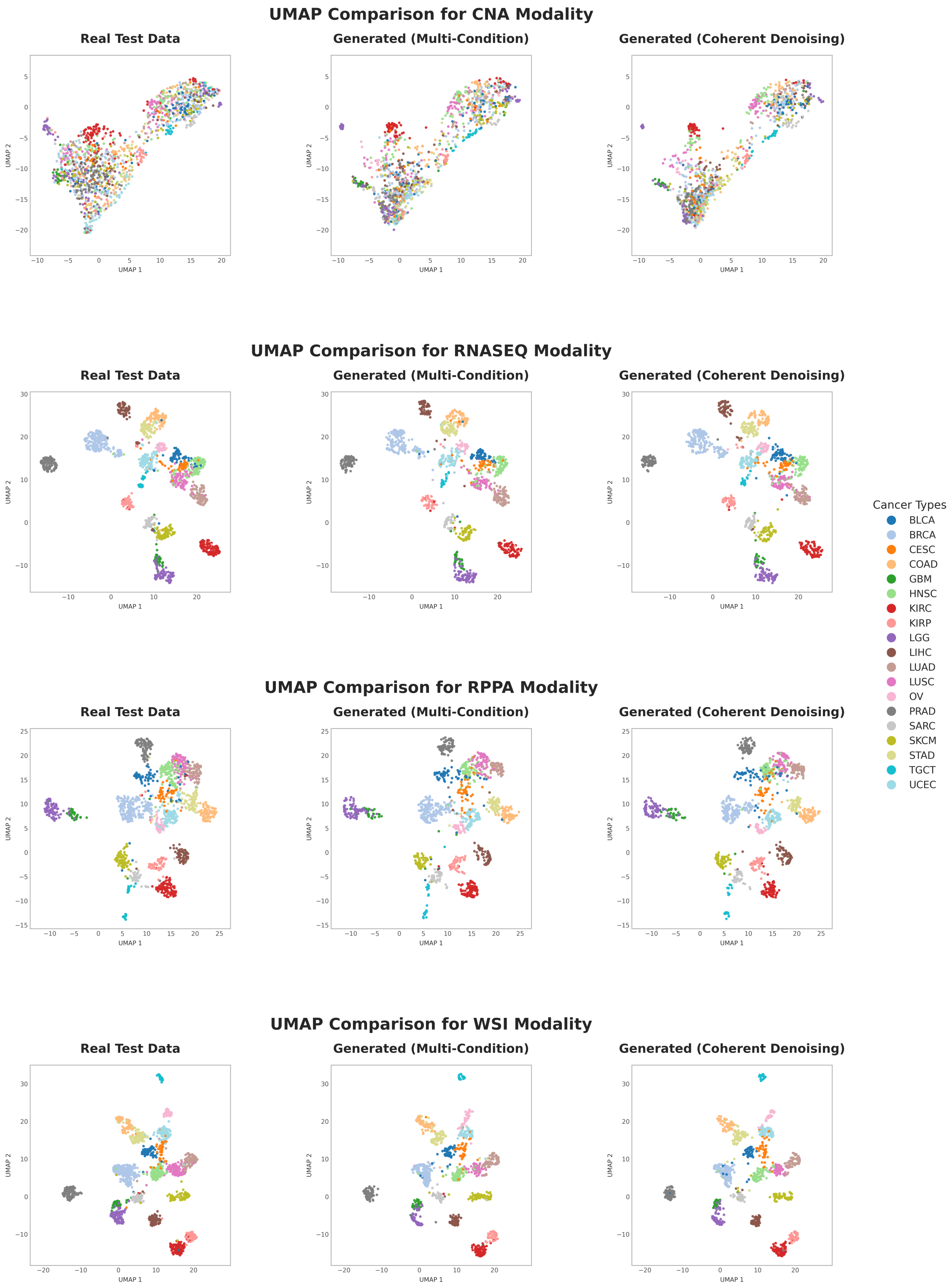


**Figure A.** Qualitative comparison of real and generated data manifolds across different data types with UMAP projections of the embeddings of each modality.

### S2 Appendix: Full Reconstruction Accuracies

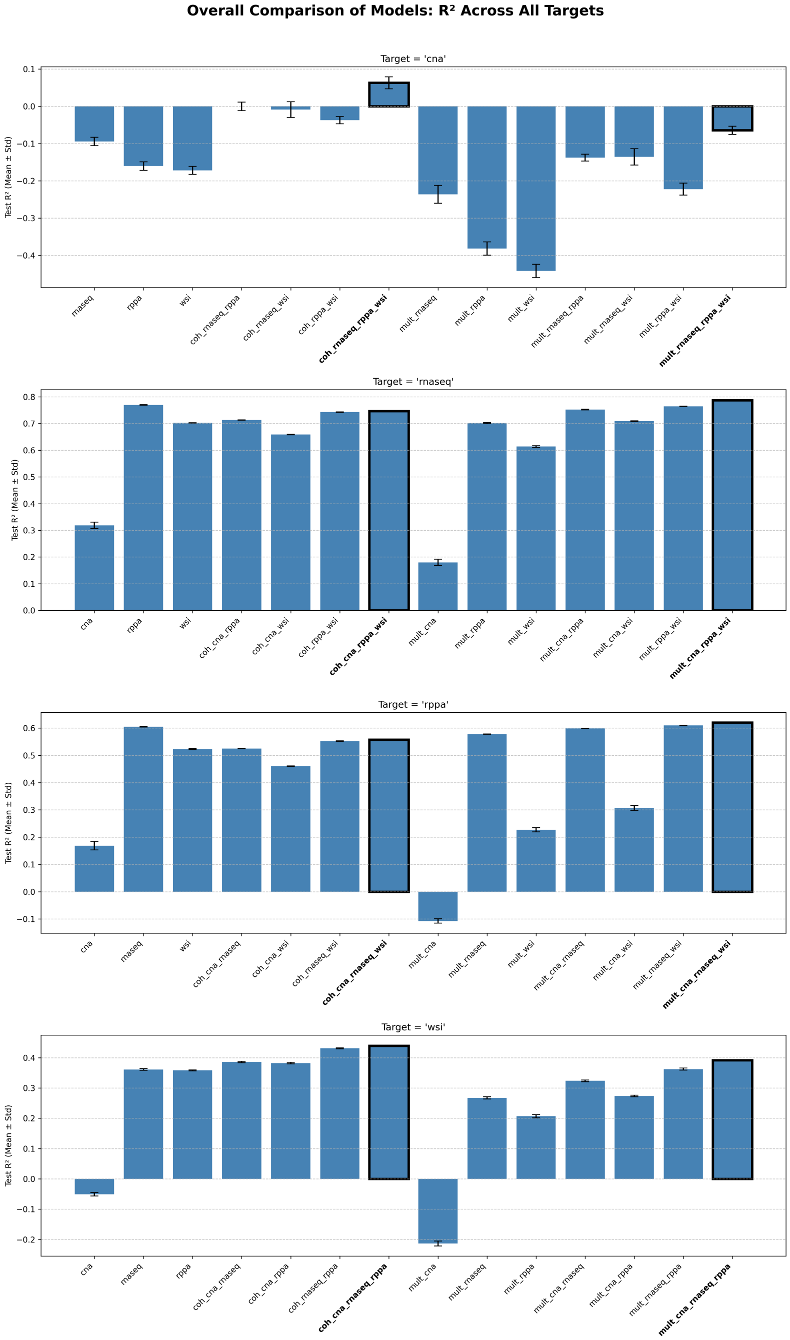


**Figure B.** Reconstruction accuracy ($R^2$) varies by target modality and generative model. Each panel displays the mean $R^2$ ($\pm$ standard deviation) from 10 generation runs for a different target modality. The x-axis compares the performance of single-condition models, the Coherent Denoising and multi-condition models, across different combinations of conditioning modalities. Highlighted bars indicate runs conditioned on all three other modalities.

| Target modality | Source modalities | Single-condition | Coherent Denoising | Multi-condition |
|---|---|---|---|---|
| **CNA** | RPPA | -0.160 ± 0.012 | — | -0.382 ± 0.018 |
| | RNASEQ | -0.094 ± 0.011 | — | -0.236 ± 0.024 |
| | WSI | -0.172 ± 0.011 | — | -0.442 ± 0.018 |
| | RPPA + WSI | — | -0.037 ± 0.010 | -0.222 ± 0.016 |
| | RNASEQ + RPPA | — | -0.000 ± 0.011 | -0.138 ± 0.009 |
| | RNASEQ + WSI | — | -0.009 ± 0.021 | -0.136 ± 0.022 |
| | RNASEQ + RPPA + WSI | — | **0.063 ± 0.016** | -0.064 ± 0.011 |
| **RNASEQ** | CNA | 0.318 ± 0.012 | — | 0.180 ± 0.012 |
| | RPPA | 0.770 ± 0.001 | — | 0.702 ± 0.002 |
| | WSI | 0.702 ± 0.001 | — | 0.614 ± 0.003 |
| | CNA + RPPA | — | 0.713 ± 0.001 | 0.752 ± 0.001 |
| | CNA + WSI | — | 0.659 ± 0.001 | 0.709 ± 0.002 |
| | RPPA + WSI | — | 0.743 ± 0.001 | 0.765 ± 0.001 |
| | CNA + RPPA + WSI | — | 0.746 ± 0.001 | **0.787 ± 0.001** |
| **RPPA** | CNA | 0.169 ± 0.016 | — | -0.107 ± 0.008 |
| | RNASEQ | 0.605 ± 0.001 | — | 0.578 ± 0.001 |
| | WSI | 0.523 ± 0.002 | — | 0.227 ± 0.008 |
| | CNA + RNASEQ | — | 0.525 ± 0.001 | 0.598 ± 0.001 |
| | CNA + WSI | — | 0.461 ± 0.001 | 0.307 ± 0.009 |
| | RNASEQ + WSI | — | 0.552 ± 0.001 | 0.610 ± 0.001 |
| | CNA + RNASEQ + WSI | — | 0.558 ± 0.001 | **0.620 ± 0.001** |
| **WSI** | CNA | -0.051 ± 0.006 | — | -0.213 ± 0.008 |
| | RPPA | 0.358 ± 0.001 | — | 0.207 ± 0.005 |
| | RNASEQ | 0.361 ± 0.003 | — | 0.267 ± 0.004 |
| | CNA + RPPA | — | 0.382 ± 0.003 | 0.274 ± 0.003 |
| | CNA + RNASEQ | — | 0.386 ± 0.002 | 0.324 ± 0.003 |
| | RNASEQ + RPPA | — | 0.431 ± 0.001 | 0.362 ± 0.003 |
| | CNA + RNASEQ + RPPA | — | **0.439 ± 0.002** | 0.392 ± 0.002 |

**Table A.** Expanded reconstruction accuracy ($R^2$) by target modality, source modalities, and generative method. Each block shows the mean $R^2$ ($\pm$ standard deviation) from 10 independent generation runs for a different target modality. Rows indicate the source modality or combination of modalities used for conditioning. Columns indicate the generative method used. The best result for each target modality is highlighted in bold.

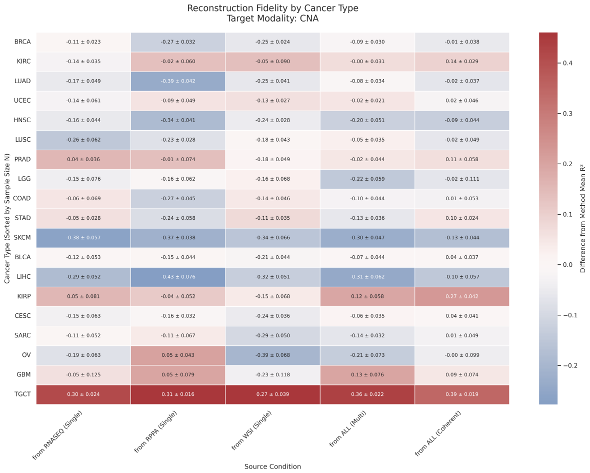


**(a)** Target Modality: CNA

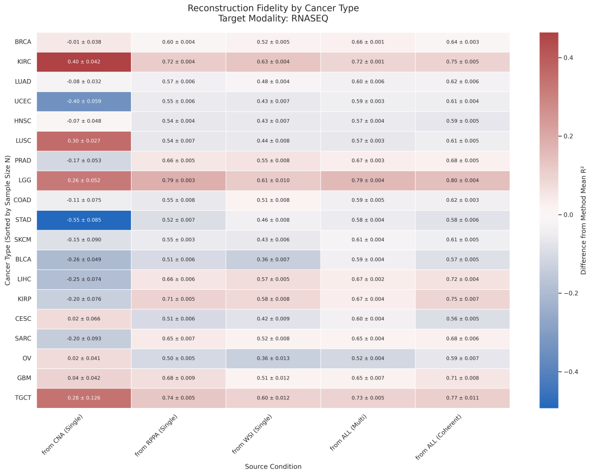


**(b)** Target Modality: RNASEQ

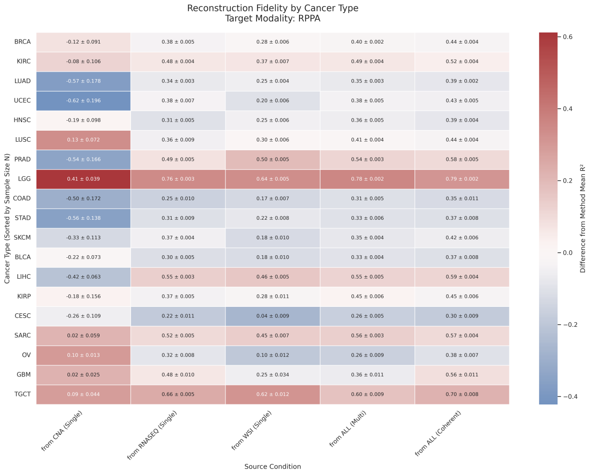


**(c)** Target Modality: RPPA

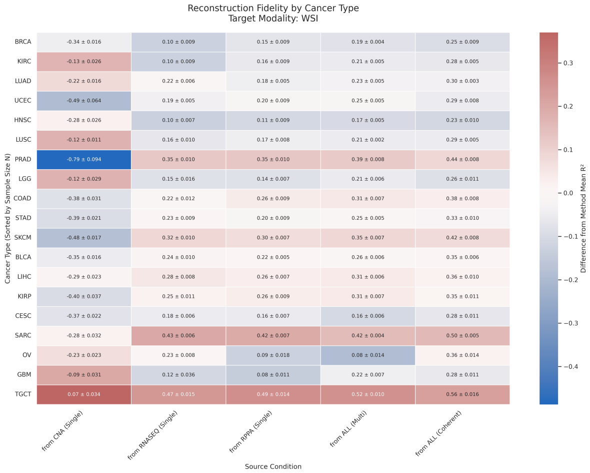


**(d)** Target Modality: WSI

**Figure C.** **Stratified Reconstruction Fidelity ($R^2$) by Cancer Type and Target Modality.** Each panel corresponds to a different target modality: (a) CNA, (b) RNASEQ, (c) RPPA, and (d) WSI. Within each panel, rows represent the 20 individual cancer types (sorted by sample size), and columns represent the source condition used for generation. The "ALL (Multi)" and "ALL (Coherent)" columns represent generation using all three other modalities as input. Cell values report the mean $R^2 \pm$ standard deviation from 10 independent generation runs. The color scale indicates the performance on a cancer type relative to the mean performance across all cancer types for that specific method.

#### S3 Appendix: Preservation of Predictive Signals in Generated Data

| Modality | Task | Data Source | Balanced Acc. | F1-Macro |
|---|---|---|---|---|
| **CNA** | Tumor Type | Real | 0.565 | 0.563 |
| | | Synthetic (Coherent) | 0.717 | 0.727 |
| | | Synthetic (Multi) | 0.647 | 0.649 |
| | Stage | Real | 0.333 | 0.318 |
| | | Synthetic (Coherent) | 0.379 | 0.369 |
| | | Synthetic (Multi) | 0.363 | 0.352 |
| **RNA-Seq** | Tumor Type | Real | 0.929 | 0.928 |
| | | Synthetic (Coherent) | 0.944 | 0.947 |
| | | Synthetic (Multi) | 0.929 | 0.931 |
| | Stage | Real | 0.489 | 0.485 |
| | | Synthetic (Coherent) | 0.498 | 0.490 |
| | | Synthetic (Multi) | 0.510 | 0.504 |
| **RPPA** | Tumor Type | Real | 0.932 | 0.901 |
| | | Synthetic (Coherent) | 0.933 | 0.935 |
| | | Synthetic (Multi) | 0.918 | 0.919 |
| | Stage | Real | 0.481 | 0.488 |
| | | Synthetic (Coherent) | 0.491 | 0.490 |
| | | Synthetic (Multi) | 0.486 | 0.485 |
| **WSI** | Tumor Type | Real | 0.937 | 0.891 |
| | | Synthetic (Coherent) | 0.938 | 0.939 |
| | | Synthetic (Multi) | 0.904 | 0.906 |
| | Stage | Real | 0.505 | 0.501 |
| | | Synthetic (Coherent) | 0.492 | 0.481 |
| | | Synthetic (Multi) | 0.499 | 0.492 |

**Table B.** Performance metrics for Random Forest classifiers trained on single real modalities and tested on either real or synthetically generated data. Synthetic data performs on-par with real data.

#### S4 Appendix: Inference-Time Data Synthesis for Downstream Predictive Tasks

| Data Condition | Experiment | Balanced Accuracy | Macro F1-Score |
|---|---|---|---|
| Full Data | Baseline | 0.528±0.006 | 0.528±0.006 |
| Cancer Label Only | Baseline | 0.366±0.026 | 0.300±0.052 |
| No CNA | Ablation | 0.518±0.011 | 0.517±0.011 |
| | Synthetic (Coherent) | 0.530±0.007 | 0.528±0.008 |
| | Synthetic (Multi) | 0.527±0.008 | 0.526±0.009 |
| No CNA, RNA-Seq | Ablation | 0.468±0.014 | 0.463±0.015 |
| | Synthetic (Coherent) | 0.524±0.007 | 0.521±0.008 |
| | Synthetic (Multi) | 0.525±0.008 | 0.520±0.008 |
| No CNA, RNA-Seq, RPPA | Ablation | 0.398±0.009 | 0.326±0.022 |
| | Synthetic (Coherent) | 0.524±0.007 | 0.514±0.007 |
| | Synthetic (Multi) | 0.515±0.009 | 0.502±0.011 |
| No CNA, RNA-Seq, WSI | Ablation | 0.392±0.028 | 0.379±0.035 |
| | Synthetic (Coherent) | 0.518±0.008 | 0.515±0.008 |
| | Synthetic (Multi) | 0.499±0.009 | 0.496±0.009 |
| No CNA, RPPA | Ablation | 0.487±0.011 | 0.462±0.014 |
| | Synthetic (Coherent) | 0.533±0.008 | 0.528±0.008 |
| | Synthetic (Multi) | 0.526±0.006 | 0.520±0.007 |
| No CNA, RPPA, WSI | Ablation | 0.473±0.010 | 0.449±0.019 |
| | Synthetic (Coherent) | 0.519±0.005 | 0.516±0.005 |
| | Synthetic (Multi) | 0.520±0.005 | 0.518±0.005 |
| No CNA, WSI | Ablation | 0.497±0.005 | 0.493±0.005 |
| | Synthetic (Coherent) | 0.522±0.007 | 0.520±0.008 |
| | Synthetic (Multi) | 0.527±0.009 | 0.525±0.009 |
| No RNA-Seq | Ablation | 0.473±0.012 | 0.483±0.010 |
| | Synthetic (Coherent) | 0.523±0.004 | 0.520±0.004 |
| | Synthetic (Multi) | 0.531±0.008 | 0.526±0.008 |
| No RNA-Seq, RPPA | Ablation | 0.450±0.019 | 0.430±0.020 |
| | Synthetic (Coherent) | 0.521±0.006 | 0.513±0.007 |
| | Synthetic (Multi) | 0.523±0.005 | 0.514±0.005 |
| No RNA-Seq, RPPA, WSI | Ablation | 0.345±0.014 | 0.304±0.012 |
| | Synthetic (Coherent) | 0.429±0.011 | 0.427±0.011 |
| | Synthetic (Multi) | 0.421±0.007 | 0.417±0.007 |
| No RNA-Seq, WSI | Ablation | 0.349±0.018 | 0.321±0.026 |
| | Synthetic (Coherent) | 0.517±0.008 | 0.514±0.008 |
| | Synthetic (Multi) | 0.511±0.011 | 0.509±0.011 |
| No RPPA | Ablation | 0.508±0.007 | 0.491±0.009 |
| | Synthetic (Coherent) | 0.527±0.008 | 0.524±0.008 |
| | Synthetic (Multi) | 0.530±0.008 | 0.526±0.007 |
| No RPPA, WSI | Ablation | 0.473±0.009 | 0.462±0.008 |
| | Synthetic (Coherent) | 0.520±0.005 | 0.517±0.007 |
| | Synthetic (Multi) | 0.522±0.007 | 0.519±0.008 |
| No WSI | Ablation | 0.497±0.008 | 0.501±0.008 |
| | Synthetic (Coherent) | 0.522±0.009 | 0.521±0.009 |
| | Synthetic (Multi) | 0.523±0.010 | 0.522±0.011 |

**Table C.** Performance metrics for Random Forest classifier (mean and standard deviation across 10 repetitions) trained on the real multimodal training set and tested on the real data with simulated missing modalities, and synthetically generated data with the Coherent Denoisng method or with the multi-condition model.

| Data Condition | Experiment Type | C-Index |
|---|---|---|
| Full Data | Baseline | 0.736±0.003 |
| Cancer Label Only | Baseline | 0.536±0.048 |
| No CNA | Ablation | 0.711±0.007 |
| | Synthetic (Coherent) | 0.739±0.002 |
| | Synthetic (Multi) | 0.737±0.003 |
| No CNA, RNA-Seq | Ablation | 0.549±0.018 |
| | Synthetic (Coherent) | 0.735±0.003 |
| | Synthetic (Multi) | 0.733±0.003 |
| No CNA, RNA-Seq, RPPA | Ablation | 0.619±0.021 |
| | Synthetic (Coherent) | 0.726±0.003 |
| | Synthetic (Multi) | 0.714±0.004 |
| No CNA, RNA-Seq, WSI | Ablation | 0.517±0.017 |
| | Synthetic (Coherent) | 0.724±0.002 |
| | Synthetic (Multi) | 0.708±0.006 |
| No CNA, RPPA | Ablation | 0.701±0.005 |
| | Synthetic (Coherent) | 0.741±0.002 |
| | Synthetic (Multi) | 0.738±0.003 |
| No CNA, RPPA, WSI | Ablation | 0.636±0.018 |
| | Synthetic (Coherent) | 0.733±0.003 |
| | Synthetic (Multi) | 0.729±0.003 |
| No CNA, WSI | Ablation | 0.654±0.009 |
| | Synthetic (Coherent) | 0.729±0.002 |
| | Synthetic (Multi) | 0.728±0.003 |
| No RNA-Seq | Ablation | 0.574±0.016 |
| | Synthetic (Coherent) | 0.734±0.004 |
| | Synthetic (Multi) | 0.736±0.004 |
| No RNA-Seq, RPPA | Ablation | 0.622±0.012 |
| | Synthetic (Coherent) | 0.730±0.004 |
| | Synthetic (Multi) | 0.729±0.004 |
| No RNA-Seq, RPPA, WSI | Ablation | 0.563±0.021 |
| | Synthetic (Coherent) | 0.664±0.006 |
| | Synthetic (Multi) | 0.652±0.006 |
| No RNA-Seq, WSI | Ablation | 0.539±0.016 |
| | Synthetic (Coherent) | 0.725±0.004 |
| | Synthetic (Multi) | 0.718±0.004 |
| No RPPA | Ablation | 0.715±0.003 |
| | Synthetic (Coherent) | 0.738±0.004 |
| | Synthetic (Multi) | 0.735±0.004 |
| No RPPA, WSI | Ablation | 0.652±0.006 |
| | Synthetic (Coherent) | 0.728±0.004 |
| | Synthetic (Multi) | 0.724±0.003 |
| No WSI | Ablation | 0.689±0.003 |
| | Synthetic (Coherent) | 0.726±0.004 |
| | Synthetic (Multi) | 0.723±0.004 |

**Table D.** C-Index for Random Survival Forest (mean and standard deviation across 10 repetitions) trained on the real multimodal training set and tested on the real data with simulated missing modalities, and synthetically generated data with the Coherent Denoisng method or with the multi-condition model.

#### S5 Appendix: Counterfactual Analysis for Survival Analysis

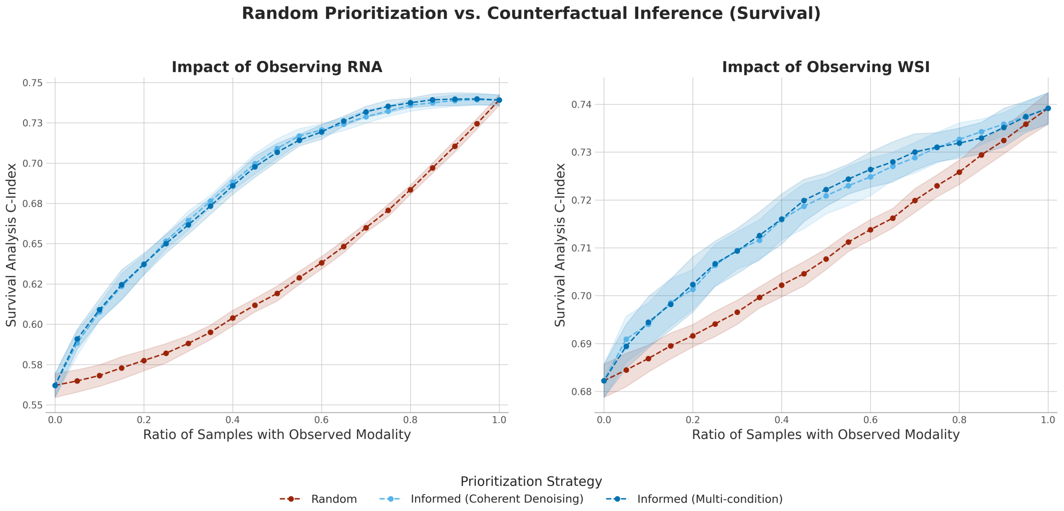


**Figure D.** Evaluating Counterfactual Inference for Prioritizing RNA-Seq (Left) and WSI (Right) Data Acquisition. The plot shows the Concordance Index of a multimodal random survival forest as the ratio of patients with an observed modality (RNA-Seq or WSI data) is varied. The Random Prioritization strategy (red) removes that modality data from patients at random. The Informed Prioritization strategies (blue) use a counterfactual variance score to preferentially acquire that modality data for the most informative patients first. Error bands show the standard deviation across 10 experimental repetitions. Note that the two plots are on separate y-axis scales, because of the intrinsic difference in performance that a predictive model has with and without that modality.

#### S6 Appendix: Statistical Tests

### 1 Preservation of Predictive Signals in Generated Data

To assess the statistical significance of performance differences between models using real vs. synthetic data, we employed a Repeated Measures ANOVA, with data modality and data source (real, coherent-generated, multi-generated) as within-subjects factors. Post-hoc analysis was conducted using paired t-tests, with p-values corrected for multiple comparisons using the Bonferroni method. A corrected alpha level was used to determine significance.

```
Repeated Measures ANOVA for: Task = tumor_type, Metric = balanced_accuracy

                    F Value   Num DF  Den DF Pr > F
modality           20256.9274 3.0000 27.0000 0.0000
data_type            736.4593 2.0000 18.0000 0.0000
modality:data_type   346.9958 6.0000 54.0000 0.0000


Repeated Measures ANOVA for: Task = tumor_type, Metric = f1_macro

                    F Value  Num DF  Den DF Pr > F
modality           6669.7201 3.0000 27.0000 0.0000
data_type           595.5192 2.0000 18.0000 0.0000
modality:data_type  126.5469 6.0000 54.0000 0.0000


Repeated Measures ANOVA for: Task = stage, Metric = balanced_accuracy

                    F Value  Num DF  Den DF Pr > F
modality           1731.9448 3.0000 27.0000 0.0000
data_type            35.8453 2.0000 18.0000 0.0000
modality:data_type   20.0905 6.0000 54.0000 0.0000


Repeated Measures ANOVA for: Task = stage, Metric = f1_macro

                    F Value  Num DF  Den DF Pr > F
modality           1367.5405 3.0000 27.0000 0.0000
data_type            13.8445 2.0000 18.0000 0.0002
modality:data_type   24.2893 6.0000 54.0000 0.0000


T-tests for the effect of data_type

ANALYSIS: Task = TUMOR_TYPE, Metric = BALANCED_ACCURACY

Modality: cna
                     real vs. synthetic_from_coherent  : p-value = 0.000000 (Significant)
                     real vs. synthetic_from_multi     : p-value = 0.000000 (Significant)
  synthetic_from_coherent vs. synthetic_from_multi     : p-value = 0.000001 (Significant)

Modality: rnaseq
                     real vs. synthetic_from_coherent  : p-value = 0.000000 (Significant)
                     real vs. synthetic_from_multi     : p-value = 0.744300 (Not Significant)
  synthetic_from_coherent vs. synthetic_from_multi     : p-value = 0.000004 (Significant)

Modality: rppa
                     real vs. synthetic_from_coherent  : p-value = 0.357651 (Not Significant)
                     real vs. synthetic_from_multi     : p-value = 0.000001 (Significant)
  synthetic_from_coherent vs. synthetic_from_multi     : p-value = 0.000000 (Significant)

Modality: wsi
                     real vs. synthetic_from_coherent  : p-value = 0.506788 (Not Significant)
                     real vs. synthetic_from_multi     : p-value = 0.000000 (Significant)
  synthetic_from_coherent vs. synthetic_from_multi     : p-value = 0.000000 (Significant)


ANALYSIS: Task = TUMOR_TYPE, Metric = F1_MACRO

Modality: cna
                     real vs. synthetic_from_coherent  : p-value = 0.000000 (Significant)
                     real vs. synthetic_from_multi     : p-value = 0.000000 (Significant)
  synthetic_from_coherent vs. synthetic_from_multi     : p-value = 0.000001 (Significant)

Modality: rnaseq
                     real vs. synthetic_from_coherent  : p-value = 0.000000 (Significant)
                     real vs. synthetic_from_multi     : p-value = 0.017115 (Not Significant)
  synthetic_from_coherent vs. synthetic_from_multi     : p-value = 0.000000 (Significant)

Modality: rppa
                     real vs. synthetic_from_coherent  : p-value = 0.001034 (Significant)
                     real vs. synthetic_from_multi     : p-value = 0.034369 (Not Significant)
```

```
  synthetic_from_coherent vs. synthetic_from_multi     : p-value = 0.000000 (Significant)

Modality: wsi
                     real vs. synthetic_from_coherent  : p-value = 0.000000 (Significant)
                     real vs. synthetic_from_multi     : p-value = 0.000001 (Significant)
  synthetic_from_coherent vs. synthetic_from_multi     : p-value = 0.000000 (Significant)


ANALYSIS: Task = STAGE, Metric = BALANCED_ACCURACY

Modality: cna
                     real vs. synthetic_from_coherent  : p-value = 0.000019 (Significant)
                     real vs. synthetic_from_multi     : p-value = 0.000083 (Significant)
  synthetic_from_coherent vs. synthetic_from_multi     : p-value = 0.021267 (Not Significant)

Modality: rnaseq
                     real vs. synthetic_from_coherent  : p-value = 0.039307 (Not Significant)
                     real vs. synthetic_from_multi     : p-value = 0.000082 (Significant)
  synthetic_from_coherent vs. synthetic_from_multi     : p-value = 0.020446 (Not Significant)

Modality: rppa
                     real vs. synthetic_from_coherent  : p-value = 0.002267 (Significant)
                     real vs. synthetic_from_multi     : p-value = 0.246810 (Not Significant)
  synthetic_from_coherent vs. synthetic_from_multi     : p-value = 0.079922 (Not Significant)

Modality: wsi
                     real vs. synthetic_from_coherent  : p-value = 0.006409 (Significant)
                     real vs. synthetic_from_multi     : p-value = 0.083884 (Not Significant)
  synthetic_from_coherent vs. synthetic_from_multi     : p-value = 0.014765 (Significant)


ANALYSIS: Task = STAGE, Metric = F1_MACRO

Modality: cna
                     real vs. synthetic_from_coherent  : p-value = 0.000061 (Significant)
                     real vs. synthetic_from_multi     : p-value = 0.000143 (Significant)
  synthetic_from_coherent vs. synthetic_from_multi     : p-value = 0.033003 (Not Significant)

Modality: rnaseq
                     real vs. synthetic_from_coherent  : p-value = 0.317641 (Not Significant)
                     real vs. synthetic_from_multi     : p-value = 0.000516 (Significant)
  synthetic_from_coherent vs. synthetic_from_multi     : p-value = 0.013981 (Significant)

Modality: rppa
                     real vs. synthetic_from_coherent  : p-value = 0.502125 (Not Significant)
                     real vs. synthetic_from_multi     : p-value = 0.502480 (Not Significant)
  synthetic_from_coherent vs. synthetic_from_multi     : p-value = 0.194758 (Not Significant)

Modality: wsi
                     real vs. synthetic_from_coherent  : p-value = 0.000575 (Significant)
                     real vs. synthetic_from_multi     : p-value = 0.034262 (Not Significant)
  synthetic_from_coherent vs. synthetic_from_multi     : p-value = 0.002183 (Significant)
```

#### 2 Inference-Time Data Synthesis for Downstream Predictive Tasks

To assess the statistical significance of the results from our downstream experiments, we employed a Repeated Measures ANOVA, with test condition and data source (ablated, coherent-generated, multi-generated) as within-subjects factors. Post-hoc analysis was conducted using paired t-tests, with p-values corrected for multiple comparisons using the Bonferroni method. A corrected alpha level was used to determine significance.

For the counterfactual analysis, we compared the overall performance of the prioritization strategies by calculating the Area Under the F1-Score Curve (AUC) for each of the 10 experimental repetitions and then performing a paired t-test on these AUC values. A corrected alpha level was used to determine significance in all post-hoc tests.

##### 2.1 Stage Classification

```
ANOVA Results for Metric: BALANCED_ACCURACY

                           F Value   Num DF   Den DF  Pr > F
test_condition            280.8313 13.0000 117.0000 0.0000
test_type                2542.4871  2.0000  18.0000 0.0000
test_condition:test_type   73.1597 26.0000 234.0000 0.0000


Post-Hoc Paired t-test Results for Metric: BALANCED_ACCURACY
Using Bonferroni-corrected alpha = 0.0012 for significance.

Condition: no_cna
          ablation vs. imputed_coherent  : p-value = 0.001249 (Not Significant)
```

```
          ablation vs. imputed_multi     : p-value = 0.071247 (Not Significant)
  imputed_coherent vs. imputed_multi     : p-value = 0.325872 (Not Significant)

Condition: no_rnaseq
          ablation vs. imputed_coherent  : p-value = 0.000000 (Significant)
          ablation vs. imputed_multi     : p-value = 0.000000 (Significant)
  imputed_coherent vs. imputed_multi     : p-value = 0.032120 (Not Significant)

Condition: no_rppa
          ablation vs. imputed_coherent  : p-value = 0.000267 (Significant)
          ablation vs. imputed_multi     : p-value = 0.000145 (Significant)
  imputed_coherent vs. imputed_multi     : p-value = 0.093417 (Not Significant)

Condition: no_wsi
          ablation vs. imputed_coherent  : p-value = 0.000011 (Significant)
          ablation vs. imputed_multi     : p-value = 0.000041 (Significant)
  imputed_coherent vs. imputed_multi     : p-value = 0.623052 (Not Significant)

Condition: no_cna_rnaseq
          ablation vs. imputed_coherent  : p-value = 0.000007 (Significant)
          ablation vs. imputed_multi     : p-value = 0.000001 (Significant)
  imputed_coherent vs. imputed_multi     : p-value = 0.787868 (Not Significant)

Condition: no_cna_rppa
          ablation vs. imputed_coherent  : p-value = 0.000001 (Significant)
          ablation vs. imputed_multi     : p-value = 0.000003 (Significant)
  imputed_coherent vs. imputed_multi     : p-value = 0.000671 (Significant)

Condition: no_cna_wsi
          ablation vs. imputed_coherent  : p-value = 0.000013 (Significant)
          ablation vs. imputed_multi     : p-value = 0.000005 (Significant)
  imputed_coherent vs. imputed_multi     : p-value = 0.166844 (Not Significant)

Condition: no_rnaseq_rppa
          ablation vs. imputed_coherent  : p-value = 0.000000 (Significant)
          ablation vs. imputed_multi     : p-value = 0.000000 (Significant)
  imputed_coherent vs. imputed_multi     : p-value = 0.562512 (Not Significant)

Condition: no_rnaseq_wsi
          ablation vs. imputed_coherent  : p-value = 0.000000 (Significant)
          ablation vs. imputed_multi     : p-value = 0.000000 (Significant)
  imputed_coherent vs. imputed_multi     : p-value = 0.099060 (Not Significant)

Condition: no_rppa_wsi
          ablation vs. imputed_coherent  : p-value = 0.000001 (Significant)
          ablation vs. imputed_multi     : p-value = 0.000001 (Significant)
  imputed_coherent vs. imputed_multi     : p-value = 0.492237 (Not Significant)

Condition: no_cna_rnaseq_rppa
          ablation vs. imputed_coherent  : p-value = 0.000000 (Significant)
          ablation vs. imputed_multi     : p-value = 0.000000 (Significant)
  imputed_coherent vs. imputed_multi     : p-value = 0.025485 (Not Significant)

Condition: no_cna_rnaseq_wsi
          ablation vs. imputed_coherent  : p-value = 0.000000 (Significant)
          ablation vs. imputed_multi     : p-value = 0.000001 (Significant)
  imputed_coherent vs. imputed_multi     : p-value = 0.000025 (Significant)

Condition: no_cna_rppa_wsi
          ablation vs. imputed_coherent  : p-value = 0.000000 (Significant)
          ablation vs. imputed_multi     : p-value = 0.000001 (Significant)
  imputed_coherent vs. imputed_multi     : p-value = 0.655039 (Not Significant)

Condition: no_rnaseq_rppa_wsi
          ablation vs. imputed_coherent  : p-value = 0.000000 (Significant)
          ablation vs. imputed_multi     : p-value = 0.000001 (Significant)
  imputed_coherent vs. imputed_multi     : p-value = 0.047397 (Not Significant)


ANOVA Results for Metric: MACRO_F1_SCORE

                           F Value   Num DF  Den DF  Pr > F
test_condition            271.6872 13.0000 117.0000 0.0000
test_type                2079.5522  2.0000  18.0000 0.0000
test_condition:test_type   84.8627 26.0000 234.0000 0.0000


Post-Hoc Paired t-test Results for Metric: MACRO_F1_SCORE
Using Bonferroni-corrected alpha = 0.0012 for significance.

Condition: no_cna
          ablation vs. imputed_coherent  : p-value = 0.004239 (Not Significant)
          ablation vs. imputed_multi     : p-value = 0.080544 (Not Significant)
```

```
  imputed_coherent vs. imputed_multi     : p-value = 0.478453 (Not Significant)

Condition: no_rnaseq
          ablation vs. imputed_coherent  : p-value = 0.000004 (Significant)
          ablation vs. imputed_multi     : p-value = 0.000000 (Significant)
  imputed_coherent vs. imputed_multi     : p-value = 0.058794 (Not Significant)

Condition: no_rppa
          ablation vs. imputed_coherent  : p-value = 0.000014 (Significant)
          ablation vs. imputed_multi     : p-value = 0.000007 (Significant)
  imputed_coherent vs. imputed_multi     : p-value = 0.195575 (Not Significant)

Condition: no_wsi
          ablation vs. imputed_coherent  : p-value = 0.000088 (Significant)
          ablation vs. imputed_multi     : p-value = 0.000509 (Significant)
  imputed_coherent vs. imputed_multi     : p-value = 0.733080 (Not Significant)

Condition: no_cna_rnaseq
          ablation vs. imputed_coherent  : p-value = 0.000013 (Significant)
          ablation vs. imputed_multi     : p-value = 0.000001 (Significant)
  imputed_coherent vs. imputed_multi     : p-value = 0.807337 (Not Significant)

Condition: no_cna_rppa
          ablation vs. imputed_coherent  : p-value = 0.000000 (Significant)
          ablation vs. imputed_multi     : p-value = 0.000000 (Significant)
  imputed_coherent vs. imputed_multi     : p-value = 0.001391 (Not Significant)

Condition: no_cna_wsi
          ablation vs. imputed_coherent  : p-value = 0.000045 (Significant)
          ablation vs. imputed_multi     : p-value = 0.000007 (Significant)
  imputed_coherent vs. imputed_multi     : p-value = 0.143245 (Not Significant)

Condition: no_rnaseq_rppa
          ablation vs. imputed_coherent  : p-value = 0.000000 (Significant)
          ablation vs. imputed_multi     : p-value = 0.000000 (Significant)
  imputed_coherent vs. imputed_multi     : p-value = 0.757921 (Not Significant)

Condition: no_rnaseq_wsi
          ablation vs. imputed_coherent  : p-value = 0.000000 (Significant)
          ablation vs. imputed_multi     : p-value = 0.000000 (Significant)
  imputed_coherent vs. imputed_multi     : p-value = 0.133177 (Not Significant)

Condition: no_rppa_wsi
          ablation vs. imputed_coherent  : p-value = 0.000000 (Significant)
          ablation vs. imputed_multi     : p-value = 0.000000 (Significant)
  imputed_coherent vs. imputed_multi     : p-value = 0.581919 (Not Significant)

Condition: no_cna_rnaseq_rppa
          ablation vs. imputed_coherent  : p-value = 0.000000 (Significant)
          ablation vs. imputed_multi     : p-value = 0.000000 (Significant)
  imputed_coherent vs. imputed_multi     : p-value = 0.012593 (Not Significant)

Condition: no_cna_rnaseq_wsi
          ablation vs. imputed_coherent  : p-value = 0.000001 (Significant)
          ablation vs. imputed_multi     : p-value = 0.000003 (Significant)
  imputed_coherent vs. imputed_multi     : p-value = 0.000021 (Significant)

Condition: no_cna_rppa_wsi
          ablation vs. imputed_coherent  : p-value = 0.000002 (Significant)
          ablation vs. imputed_multi     : p-value = 0.000002 (Significant)
  imputed_coherent vs. imputed_multi     : p-value = 0.557439 (Not Significant)

Condition: no_rnaseq_rppa_wsi
          ablation vs. imputed_coherent  : p-value = 0.000000 (Significant)
          ablation vs. imputed_multi     : p-value = 0.000000 (Significant)
  imputed_coherent vs. imputed_multi     : p-value = 0.024217 (Not Significant)


ANALYSIS: Imputation vs. Full Data for Metric: BALANCED_ACCURACY
Using Bonferroni-corrected alpha = 0.0018 for significance.

Condition: no_cna (vs. Full Data)
  imputed_coherent : p-value = 0.619192 (Not Significant)
     imputed_multi : p-value = 0.605773 (Not Significant)

Condition: no_rnaseq (vs. Full Data)
  imputed_coherent : p-value = 0.016879 (Not Significant)
     imputed_multi : p-value = 0.536730 (Not Significant)

Condition: no_rppa (vs. Full Data)
  imputed_coherent : p-value = 0.525017 (Not Significant)
     imputed_multi : p-value = 0.675987 (Not Significant)
```

```
Condition: no_wsi (vs. Full Data)
  imputed_coherent : p-value = 0.032724 (Not Significant)
     imputed_multi : p-value = 0.151735 (Not Significant)

Condition: no_cna_rnaseq (vs. Full Data)
  imputed_coherent : p-value = 0.126656 (Not Significant)
     imputed_multi : p-value = 0.396201 (Not Significant)

Condition: no_cna_rppa (vs. Full Data)
  imputed_coherent : p-value = 0.183691 (Not Significant)
     imputed_multi : p-value = 0.313760 (Not Significant)

Condition: no_cna_wsi (vs. Full Data)
  imputed_coherent : p-value = 0.047027 (Not Significant)
     imputed_multi : p-value = 0.565660 (Not Significant)

Condition: no_rnaseq_rppa (vs. Full Data)
  imputed_coherent : p-value = 0.039394 (Not Significant)
     imputed_multi : p-value = 0.079447 (Not Significant)

Condition: no_rnaseq_wsi (vs. Full Data)
  imputed_coherent : p-value = 0.000271 (Significant (full_data is better))
     imputed_multi : p-value = 0.000121 (Significant (full_data is better))

Condition: no_rppa_wsi (vs. Full Data)
  imputed_coherent : p-value = 0.013246 (Not Significant)
     imputed_multi : p-value = 0.006305 (Not Significant)

Condition: no_cna_rnaseq_rppa (vs. Full Data)
  imputed_coherent : p-value = 0.137558 (Not Significant)
     imputed_multi : p-value = 0.010216 (Not Significant)

Condition: no_cna_rnaseq_wsi (vs. Full Data)
  imputed_coherent : p-value = 0.000504 (Significant (full_data is better))
     imputed_multi : p-value = 0.000002 (Significant (full_data is better))

Condition: no_cna_rppa_wsi (vs. Full Data)
  imputed_coherent : p-value = 0.000538 (Significant (full_data is better))
     imputed_multi : p-value = 0.005403 (Not Significant)

Condition: no_rnaseq_rppa_wsi (vs. Full Data)
  imputed_coherent : p-value = 0.000000 (Significant (full_data is better))
     imputed_multi : p-value = 0.000000 (Significant (full_data is better))


ANALYSIS: Imputation vs. Full Data for Metric: MACRO_F1_SCORE
Using Bonferroni-corrected alpha = 0.0018 for significance.

Condition: no_cna (vs. Full Data)
  imputed_coherent : p-value = 0.970809 (Not Significant)
     imputed_multi : p-value = 0.515542 (Not Significant)

Condition: no_rnaseq (vs. Full Data)
  imputed_coherent : p-value = 0.002692 (Not Significant)
     imputed_multi : p-value = 0.656059 (Not Significant)

Condition: no_rppa (vs. Full Data)
  imputed_coherent : p-value = 0.176310 (Not Significant)
     imputed_multi : p-value = 0.555354 (Not Significant)

Condition: no_wsi (vs. Full Data)
  imputed_coherent : p-value = 0.013130 (Not Significant)
     imputed_multi : p-value = 0.122376 (Not Significant)

Condition: no_cna_rnaseq (vs. Full Data)
  imputed_coherent : p-value = 0.040785 (Not Significant)
     imputed_multi : p-value = 0.083705 (Not Significant)

Condition: no_cna_rppa (vs. Full Data)
  imputed_coherent : p-value = 0.922261 (Not Significant)
     imputed_multi : p-value = 0.016928 (Not Significant)

Condition: no_cna_wsi (vs. Full Data)
  imputed_coherent : p-value = 0.021443 (Not Significant)
     imputed_multi : p-value = 0.281746 (Not Significant)

Condition: no_rnaseq_rppa (vs. Full Data)
  imputed_coherent : p-value = 0.000937 (Significant (full_data is better))
     imputed_multi : p-value = 0.000780 (Significant (full_data is better))

Condition: no_rnaseq_wsi (vs. Full Data)
  imputed_coherent : p-value = 0.000202 (Significant (full_data is better))
     imputed_multi : p-value = 0.000098 (Significant (full_data is better))
```

```
Condition: no_rppa_wsi (vs. Full Data)
  imputed_coherent : p-value = 0.007403 (Not Significant)
     imputed_multi : p-value = 0.002582 (Not Significant)

Condition: no_cna_rnaseq_rppa (vs. Full Data)
  imputed_coherent : p-value = 0.001814 (Not Significant)
     imputed_multi : p-value = 0.000429 (Significant (full_data is better))

Condition: no_cna_rnaseq_wsi (vs. Full Data)
  imputed_coherent : p-value = 0.000191 (Significant (full_data is better))
     imputed_multi : p-value = 0.000000 (Significant (full_data is better))

Condition: no_cna_rppa_wsi (vs. Full Data)
  imputed_coherent : p-value = 0.000559 (Significant (full_data is better))
     imputed_multi : p-value = 0.001492 (Significant (full_data is better))

Condition: no_rnaseq_rppa_wsi (vs. Full Data)
  imputed_coherent : p-value = 0.000000 (Significant (full_data is better))
     imputed_multi : p-value = 0.000000 (Significant (full_data is better))
```

#### 2.2 Survival Analysis

```
ANOVA Results for Metric: C_INDEX

                           F Value   Num DF  Den DF  Pr > F
test_condition            722.5935 13.0000 117.0000 0.0000
test_type                1968.5437  4.0000  36.0000 0.0000
test_condition:test_type  154.8086 52.0000 468.0000 0.0000


Post-Hoc Paired t-test Results for Metric: C_INDEX
Using Bonferroni-corrected alpha = 0.0012 for significance.

Condition: no_cna
          ablation vs. imputed_coherent  : p-value = 0.000000 (Significant)
          ablation vs. imputed_multi     : p-value = 0.000001 (Significant)
  imputed_coherent vs. imputed_multi     : p-value = 0.063451 (Not Significant)

Condition: no_rnaseq
          ablation vs. imputed_coherent  : p-value = 0.000000 (Significant)
          ablation vs. imputed_multi     : p-value = 0.000000 (Significant)
  imputed_coherent vs. imputed_multi     : p-value = 0.026032 (Not Significant)

Condition: no_rppa
          ablation vs. imputed_coherent  : p-value = 0.000000 (Significant)
          ablation vs. imputed_multi     : p-value = 0.000000 (Significant)
  imputed_coherent vs. imputed_multi     : p-value = 0.001847 (Not Significant)

Condition: no_wsi
          ablation vs. imputed_coherent  : p-value = 0.000000 (Significant)
          ablation vs. imputed_multi     : p-value = 0.000000 (Significant)
  imputed_coherent vs. imputed_multi     : p-value = 0.027150 (Not Significant)

Condition: no_cna_rnaseq
          ablation vs. imputed_coherent  : p-value = 0.000000 (Significant)
          ablation vs. imputed_multi     : p-value = 0.000000 (Significant)
  imputed_coherent vs. imputed_multi     : p-value = 0.163051 (Not Significant)

Condition: no_cna_rppa
          ablation vs. imputed_coherent  : p-value = 0.000000 (Significant)
          ablation vs. imputed_multi     : p-value = 0.000000 (Significant)
  imputed_coherent vs. imputed_multi     : p-value = 0.003526 (Not Significant)

Condition: no_cna_wsi
          ablation vs. imputed_coherent  : p-value = 0.000000 (Significant)
          ablation vs. imputed_multi     : p-value = 0.000000 (Significant)
  imputed_coherent vs. imputed_multi     : p-value = 0.530708 (Not Significant)

Condition: no_rnaseq_rppa
          ablation vs. imputed_coherent  : p-value = 0.000000 (Significant)
          ablation vs. imputed_multi     : p-value = 0.000000 (Significant)
  imputed_coherent vs. imputed_multi     : p-value = 0.095726 (Not Significant)

Condition: no_rnaseq_wsi
          ablation vs. imputed_coherent  : p-value = 0.000000 (Significant)
          ablation vs. imputed_multi     : p-value = 0.000000 (Significant)
  imputed_coherent vs. imputed_multi     : p-value = 0.001713 (Not Significant)

Condition: no_rppa_wsi
          ablation vs. imputed_coherent  : p-value = 0.000000 (Significant)
          ablation vs. imputed_multi     : p-value = 0.000000 (Significant)
  imputed_coherent vs. imputed_multi     : p-value = 0.005730 (Not Significant)
```

```
Condition: no_cna_rnaseq_rppa
          ablation vs. imputed_coherent  : p-value = 0.000000 (Significant)
          ablation vs. imputed_multi     : p-value = 0.000000 (Significant)
  imputed_coherent vs. imputed_multi     : p-value = 0.000060 (Significant)

Condition: no_cna_rnaseq_wsi
          ablation vs. imputed_coherent  : p-value = 0.000000 (Significant)
          ablation vs. imputed_multi     : p-value = 0.000000 (Significant)
  imputed_coherent vs. imputed_multi     : p-value = 0.000020 (Significant)

Condition: no_cna_rppa_wsi
          ablation vs. imputed_coherent  : p-value = 0.000000 (Significant)
          ablation vs. imputed_multi     : p-value = 0.000000 (Significant)
  imputed_coherent vs. imputed_multi     : p-value = 0.021486 (Not Significant)

Condition: no_rnaseq_rppa_wsi
          ablation vs. imputed_coherent  : p-value = 0.000000 (Significant)
          ablation vs. imputed_multi     : p-value = 0.000000 (Significant)
  imputed_coherent vs. imputed_multi     : p-value = 0.003909 (Not Significant)


ANALYSIS: Imputation vs. Full Data for Metric: C_INDEX
Using Bonferroni-corrected alpha = 0.0018 for significance.

Condition: no_cna (vs. Full Data)
  imputed_coherent : p-value = 0.008620 (Not Significant)
     imputed_multi : p-value = 0.322543 (Not Significant)

Condition: no_rnaseq (vs. Full Data)
  imputed_coherent : p-value = 0.021880 (Not Significant)
     imputed_multi : p-value = 0.885021 (Not Significant)

Condition: no_rppa (vs. Full Data)
  imputed_coherent : p-value = 0.008366 (Not Significant)
     imputed_multi : p-value = 0.100889 (Not Significant)

Condition: no_wsi (vs. Full Data)
  imputed_coherent : p-value = 0.000000 (Significant (full_data is better))
     imputed_multi : p-value = 0.000001 (Significant (full_data is better))

Condition: no_cna_rnaseq (vs. Full Data)
  imputed_coherent : p-value = 0.497254 (Not Significant)
     imputed_multi : p-value = 0.025909 (Not Significant)

Condition: no_cna_rppa (vs. Full Data)
  imputed_coherent : p-value = 0.000585 (Significant (imputed_coherent is better))
     imputed_multi : p-value = 0.221189 (Not Significant)

Condition: no_cna_wsi (vs. Full Data)
  imputed_coherent : p-value = 0.001057 (Significant (full_data is better))
     imputed_multi : p-value = 0.000103 (Significant (full_data is better))

Condition: no_rnaseq_rppa (vs. Full Data)
  imputed_coherent : p-value = 0.000046 (Significant (full_data is better))
     imputed_multi : p-value = 0.000014 (Significant (full_data is better))

Condition: no_rnaseq_wsi (vs. Full Data)
  imputed_coherent : p-value = 0.000003 (Significant (full_data is better))
     imputed_multi : p-value = 0.000001 (Significant (full_data is better))

Condition: no_rppa_wsi (vs. Full Data)
  imputed_coherent : p-value = 0.000941 (Significant (full_data is better))
     imputed_multi : p-value = 0.000001 (Significant (full_data is better))

Condition: no_cna_rnaseq_rppa (vs. Full Data)
  imputed_coherent : p-value = 0.000037 (Significant (full_data is better))
     imputed_multi : p-value = 0.000000 (Significant (full_data is better))

Condition: no_cna_rnaseq_wsi (vs. Full Data)
  imputed_coherent : p-value = 0.000000 (Significant (full_data is better))
     imputed_multi : p-value = 0.000000 (Significant (full_data is better))

Condition: no_cna_rppa_wsi (vs. Full Data)
  imputed_coherent : p-value = 0.086749 (Not Significant)
     imputed_multi : p-value = 0.001257 (Significant (full_data is better))

Condition: no_rnaseq_rppa_wsi (vs. Full Data)
  imputed_coherent : p-value = 0.000000 (Significant (full_data is better))
     imputed_multi : p-value = 0.000000 (Significant (full_data is better))
```

#### 3 Counterfactual Analysis

```
Statistical Analysis: Counterfactual Inference vs. Random Ablation
Metric: Area Under the F1-Score Curve (AUC)
Using Bonferroni-corrected alpha = 0.0167 for significance.

Ablated Modality: RNA
    random vs. coherent  : p-value = 0.000000 (Significant (coherent is better))
    random vs. multi     : p-value = 0.000000 (Significant (multi is better))
  coherent vs. multi     : p-value = 0.626752 (Not Significant)

Ablated Modality: WSI
    random vs. coherent  : p-value = 0.000000 (Significant (coherent is better))
    random vs. multi     : p-value = 0.000000 (Significant (multi is better))
  coherent vs. multi     : p-value = 0.057699 (Not Significant)
```

#### S7 Appendix: Hyperparameters

| Model (Target/Condition) | batch_size | learning_rate | initial_size | n_layers | time_embedding_dimension | cond_embedding_dim |
|---|---|---|---|---|---|---|
| cna_from_rnaseq | 64 | 0.0001 | 1024 | 5 | 128 | 32 |
| cna_from_rppa | 128 | 0.0001 | 1024 | 5 | 128 | 32 |
| cna_from_wsi | 128 | 0.0001 | 1024 | 5 | 128 | 32 |
| cna_from_multi | 128 | 0.0001 | 1024 | 4 | 128 | 32 |
| rnaseq_from_cna | 128 | 0.0001 | 1024 | 5 | 64 | 32 |
| rnaseq_from_rppa | 128 | 0.0001 | 1024 | 5 | 64 | 32 |
| rnaseq_from_wsi | 64 | 0.0001 | 1024 | 7 | 64 | 32 |
| rnaseq_from_multi | 128 | 0.0001 | 1024 | 5 | 64 | 32 |
| rppa_from_cna | 128 | 0.0001 | 1024 | 6 | 64 | 32 |
| rppa_from_rnaseq | 128 | 0.0001 | 1024 | 7 | 128 | 32 |
| rppa_from_wsi | 128 | 0.0001 | 1024 | 6 | 64 | 32 |
| rppa_from_multi | 128 | 0.0001 | 1024 | 6 | 128 | 32 |
| wsi_from_cna | 128 | 0.0001 | 1024 | 6 | 64 | 32 |
| wsi_from_rnaseq | 128 | 0.0001 | 1024 | 5 | 64 | 32 |
| wsi_from_rppa | 128 | 0.0001 | 1024 | 6 | 128 | 32 |
| wsi_from_multi | 128 | 0.0001 | 1024 | 7 | 64 | 32 |

**Table E.** Best hyperparmeters for each trained model

#### S8 Appendix: Coherent Denoising Ensemble Weights

This appendix details the performance-based weighting scheme used in the Coherent Denoising ensemble. The weights are assigned to prioritize high-fidelity modalities while down-weighting noisier inputs, ensuring the generation process is driven by the most reliable available signal.

| Target Modality | Source Modality | Validation Loss (MSE) |
|---|---|---|
| **CNA** | RNA-Seq | 1.006 |
| | RPPA | 1.096 |
| | WSI | 1.086 |
| **RNA-Seq** | CNA | 0.640 |
| | RPPA | 0.220 |
| | WSI | 0.294 |
| **RPPA** | CNA | 0.778 |
| | RNA-Seq | 0.367 |
| | WSI | 0.437 |
| **WSI** | CNA | 1.049 |
| | RNA-Seq | 0.642 |
| | RPPA | 0.636 |

**Table F.** Validation Mean Squared Error (MSE) per Conditioning Pair. The weights ($w_i$) for the ensemble are calculated as the Softmax of the inverse validation MSE ($1/L$) for each available single-condition model. The table below lists the raw MSE values ($L$) used for this calculation. Lower MSE indicates higher reconstruction fidelity for that specific source-target pair.

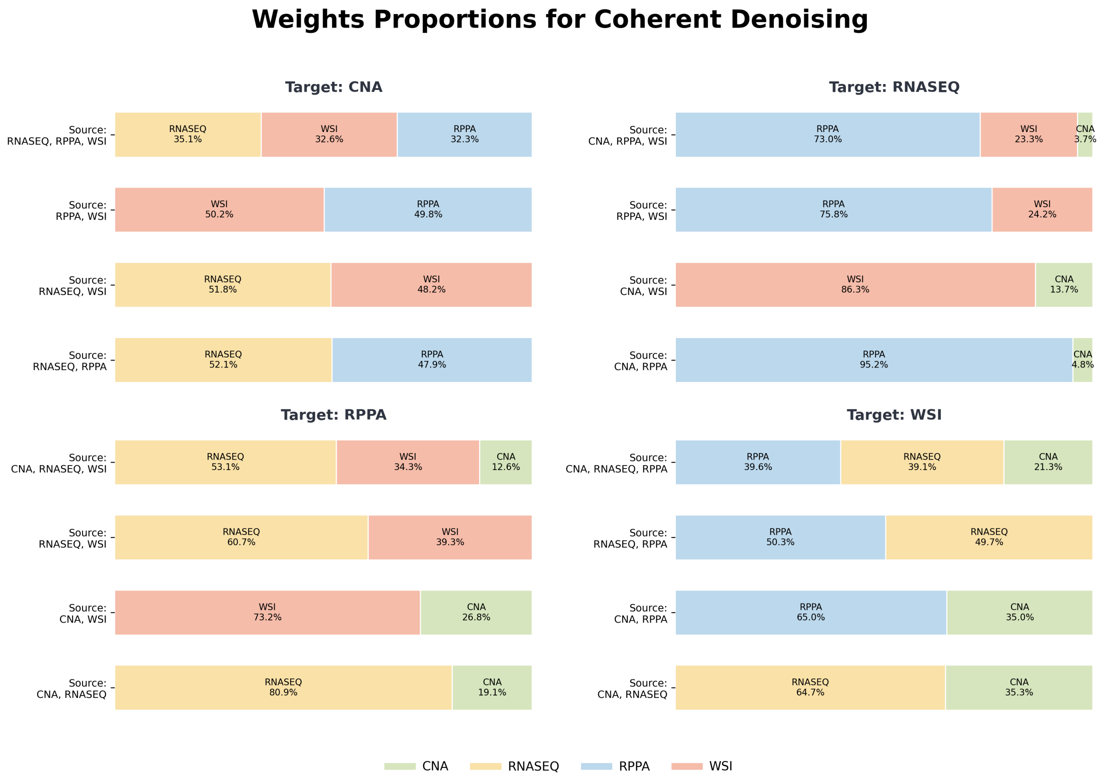


**Figure E.** Modality-Specific Influence in the Coherent Denoising Ensemble. Each panel illustrates the relative contribution (consensus share) of available conditioning modalities toward the generation of a specific target modality (CNA, RNA-Seq, RPPA, or WSI). Within each stacked bar, the segments are ordered from the highest influence to the lowest , reflecting the hierarchical prioritization of the models.

#### S9 Appendix: Sensitivity Analysis of Rejection Sampling

This appendix presents a sensitivity analysis of the Coherent Denoising rejection sampling mechanism. To evaluate the robustness of the framework, we show the effect of varying the cosine distance threshold ($\tau_{cos}$) across a range of ±30% from the default value of 1.0.

| Cosine Threshold | cna | rnaseq | rppa | wsi |
|---|---|---|---|---|
| **0.7 (-30%)** | 21.81% ± 0.69 | 7.74% ± 0.79 | 16.38% ± 0.57 | 13.22% ± 0.42 |
| **0.8 (-20%)** | 8.61% ± 0.59 | 2.69% ± 0.37 | 5.76% ± 0.37 | 4.07% ± 0.56 |
| **0.9 (-10%)** | 2.44% ± 0.47 | 0.93% ± 0.17 | 1.76% ± 0.31 | 1.10% ± 0.16 |
| **1.0 (default)** | 0.32% ± 0.16 | 0.24% ± 0.11 | 0.44% ± 0.14 | 0.26% ± 0.08 |
| **1.1 (+10%)** | 0.02% ± 0.03 | 0.07% ± 0.07 | 0.03% ± 0.04 | 0.05% ± 0.06 |
| **1.2 (+20%)** | 0.00% ± 0.00 | 0.00% ± 0.00 | 0.01% ± 0.02 | 0.00% ± 0.00 |
| **1.3 (+30%)** | 0.00% ± 0.00 | 0.00% ± 0.00 | 0.00% ± 0.00 | 0.00% ± 0.00 |

**Table G.** Impact of Cosine Threshold on Rejection Rate. The table reports the mean Rejection Rate ± standard deviation across 10 independent experimental runs.

| Cosine Threshold | cna | rnaseq | rppa | wsi |
|---|---|---|---|---|
| **0.7 (-30%)** | 0.0422 ± 0.0183 | 0.7975 ± 0.0005 | 0.6433 ± 0.0010 | 0.4524 ± 0.0015 |
| **0.8 (-20%)** | 0.0518 ± 0.0128 | 0.7978 ± 0.0007 | 0.6441 ± 0.0009 | 0.4520 ± 0.0014 |
| **0.9 (-10%)** | 0.0453 ± 0.0126 | 0.7978 ± 0.0006 | 0.6439 ± 0.0011 | 0.4526 ± 0.0008 |
| **1.0 (default)** | 0.0407 ± 0.0052 | 0.7980 ± 0.0004 | 0.6443 ± 0.0013 | 0.4524 ± 0.0019 |
| **1.1 (+10%)** | 0.0464 ± 0.0090 | 0.7977 ± 0.0007 | 0.6442 ± 0.0009 | 0.4527 ± 0.0017 |
| **1.2 (+20%)** | 0.0516 ± 0.0153 | 0.7978 ± 0.0005 | 0.6444 ± 0.0012 | 0.4530 ± 0.0017 |
| **1.3 (+30%)** | 0.0469 ± 0.0147 | 0.7978 ± 0.0004 | 0.6437 ± 0.0011 | 0.4520 ± 0.0021 |

**Table H.** Impact of Cosine Threshold on Reconstruction Fidelity. The table reports the mean Reconstruction Error ($R^2$) ± standard deviation across 10 independent experimental runs. To provide an estimate of stability, $R^2$ scores were calculated on the full generated cohort, including samples that would have been rejected at that specific threshold.

#### S10 Appendix: Impact of Embedding Dimension

In this appendix we compare the default latent dimension (32) against larger dimensions (64, 128, 256). This analysis evaluates both the baseline predictive performance of the real data and the generative capability of the framework in higher-dimensional spaces.

##### UMAP

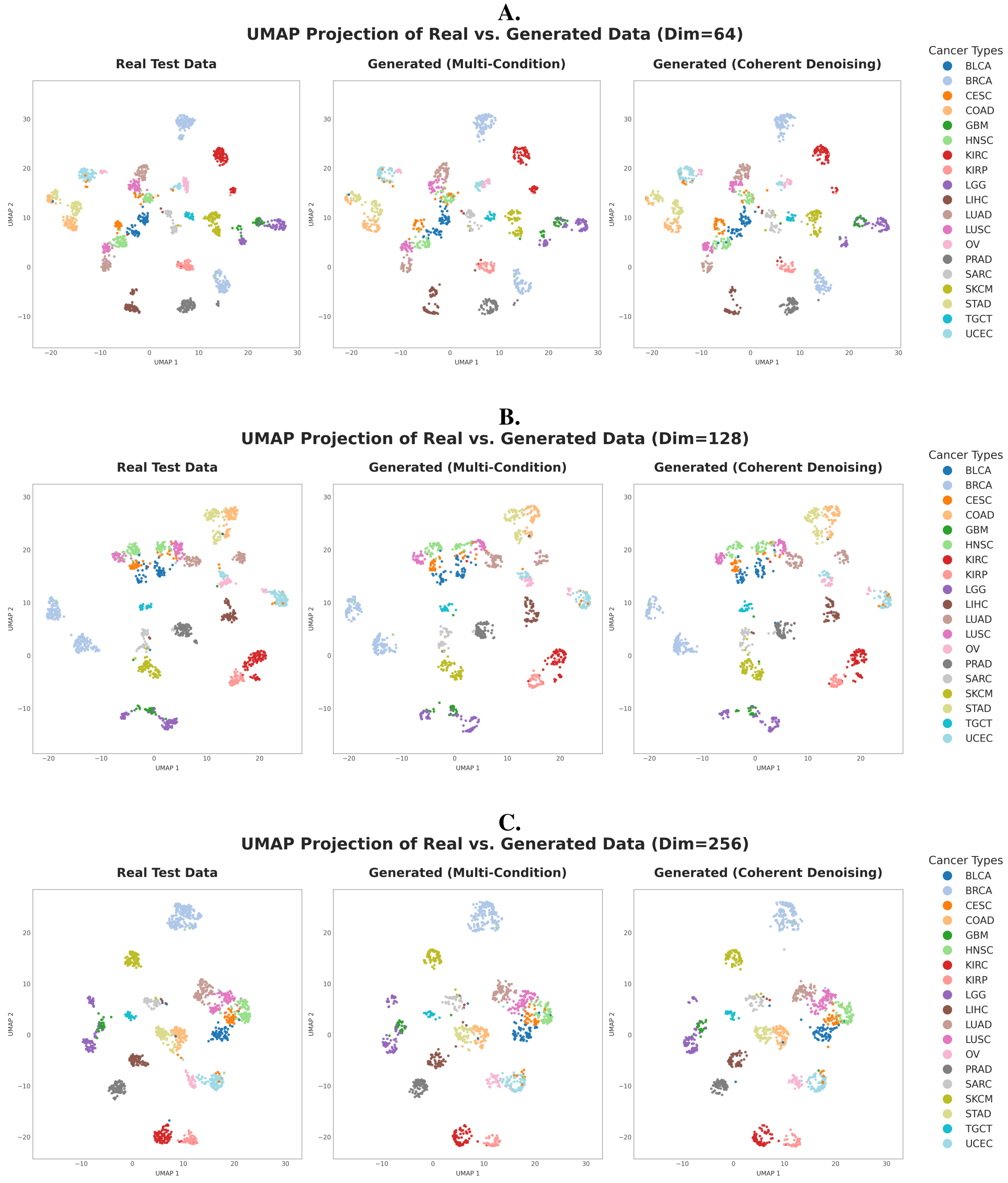


**Figure F. UMAP projections across Latent Dimensions.** UMAP projections of multimodal embeddings for dimensions 64 (A), 128 (B), and 256 (C). In each row, the ground-truth data (Left) is compared with reconstructions from the Multi-condition model (Middle) and the Coherent Denoising ensemble (Right). The visualization confirms that the generative framework maintains structural integrity and distinct cancer-type clustering even as the dimensionality of the latent space increases.

##### Downstream Tasks

| Embedding Dim | Stage Classification (F1 Score) | Survival Analysis (C-Index) |
|---|---|---|
| **32 (Default)** | **0.528 ± 0.006** | **0.736 ± 0.003** |
| **64** | 0.519 ± 0.008 | 0.732 ± 0.004 |
| **128** | 0.511 ± 0.008 | 0.734 ± 0.004 |
| **256** | 0.513 ± 0.014 | 0.727 ± 0.003 |

**Table I.** Baseline Performance of Real Data across Latent Dimensions. We evaluated the predictive performance of Multimodal Random Forest classifiers and Random Survival Forests trained on the real data, and tested on ground-truth embeddings at varying dimensions. Values represent Mean ± Standard Deviation across 10 experimental runs.

| **Embedding Dim** | **Test Condition** | **Stage Classification (F1 Score)** | | | **Survival Analysis (C-Index)** | | |
|---|---|---|---|---|---|---|---|
| | | **Ablation** | **Gain from generated data** | | **Ablation** | **Gain from generated data** | |
| | | **Drop** | **Multi-cond.** | **Coherent** | **Drop** | **Multi-cond.** | **Coherent** |
| **64** | No Cna | -0.011 ± 0.014 | +0.007 ± 0.015 | +0.012 ± 0.014 | -0.030 ± 0.005 | +0.032 ± 0.006 | +0.031 ± 0.006 |
| | No Rnaseq | -0.056 ± 0.023 | +0.058 ± 0.017 | +0.055 ± 0.018 | -0.166 ± 0.006 | +0.167 ± 0.008 | +0.167 ± 0.008 |
| | No Rppa | -0.046 ± 0.010 | +0.046 ± 0.016 | +0.050 ± 0.014 | -0.024 ± 0.004 | +0.028 ± 0.003 | +0.026 ± 0.003 |
| | No Wsi | -0.030 ± 0.011 | +0.031 ± 0.011 | +0.031 ± 0.013 | -0.048 ± 0.005 | +0.038 ± 0.005 | +0.044 ± 0.006 |
| | No Cna, Rnaseq | -0.062 ± 0.024 | +0.050 ± 0.023 | +0.062 ± 0.021 | -0.179 ± 0.016 | +0.177 ± 0.017 | +0.182 ± 0.015 |
| | No Cna, Rppa | -0.067 ± 0.013 | +0.065 ± 0.011 | +0.068 ± 0.013 | -0.048 ± 0.009 | +0.051 ± 0.010 | +0.050 ± 0.011 |
| | No Cna, Wsi | -0.033 ± 0.010 | +0.035 ± 0.014 | +0.031 ± 0.014 | -0.080 ± 0.008 | +0.073 ± 0.009 | +0.078 ± 0.008 |
| | No Rnaseq, Rppa | -0.085 ± 0.029 | +0.068 ± 0.026 | +0.073 ± 0.029 | -0.147 ± 0.008 | +0.146 ± 0.010 | +0.146 ± 0.009 |
| | No Rnaseq, Wsi | -0.243 ± 0.032 | +0.232 ± 0.034 | +0.237 ± 0.035 | -0.172 ± 0.013 | +0.162 ± 0.014 | +0.166 ± 0.013 |
| | No Rppa, Wsi | -0.065 ± 0.008 | +0.065 ± 0.008 | +0.062 ± 0.009 | -0.087 ± 0.007 | +0.079 ± 0.007 | +0.083 ± 0.007 |
| | No Cna, Rnaseq, Rppa | -0.156 ± 0.035 | +0.136 ± 0.034 | +0.147 ± 0.038 | -0.163 ± 0.007 | +0.158 ± 0.007 | +0.162 ± 0.009 |
| | No Cna, Rnaseq, Wsi | -0.229 ± 0.039 | +0.200 ± 0.044 | +0.225 ± 0.039 | -0.184 ± 0.015 | +0.168 ± 0.011 | +0.180 ± 0.013 |
| | No Cna, Rppa, Wsi | -0.081 ± 0.015 | +0.081 ± 0.015 | +0.076 ± 0.016 | -0.114 ± 0.015 | +0.106 ± 0.015 | +0.112 ± 0.015 |
| | No Rnaseq, Rppa, Wsi | -0.260 ± 0.035 | +0.144 ± 0.034 | +0.160 ± 0.030 | -0.168 ± 0.018 | +0.094 ± 0.024 | +0.111 ± 0.021 |
| **128** | No Cna | -0.017 ± 0.011 | +0.022 ± 0.010 | +0.020 ± 0.010 | -0.039 ± 0.010 | +0.038 ± 0.009 | +0.037 ± 0.010 |
| | No Rnaseq | -0.084 ± 0.019 | +0.090 ± 0.020 | +0.089 ± 0.022 | -0.184 ± 0.006 | +0.182 ± 0.008 | +0.185 ± 0.007 |
| | No Rppa | -0.037 ± 0.011 | +0.042 ± 0.013 | +0.039 ± 0.012 | -0.024 ± 0.007 | +0.027 ± 0.006 | +0.026 ± 0.006 |
| | No Wsi | -0.027 ± 0.010 | +0.031 ± 0.009 | +0.033 ± 0.008 | -0.038 ± 0.010 | +0.029 ± 0.007 | +0.031 ± 0.008 |
| | No Cna, Rnaseq | -0.094 ± 0.035 | +0.094 ± 0.030 | +0.099 ± 0.032 | -0.197 ± 0.013 | +0.189 ± 0.011 | +0.196 ± 0.012 |
| | No Cna, Rppa | -0.066 ± 0.008 | +0.061 ± 0.011 | +0.064 ± 0.010 | -0.055 ± 0.011 | +0.059 ± 0.012 | +0.056 ± 0.010 |
| | No Cna, Wsi | -0.042 ± 0.011 | +0.048 ± 0.009 | +0.046 ± 0.010 | -0.091 ± 0.008 | +0.082 ± 0.008 | +0.081 ± 0.007 |
| | No Rnaseq, Rppa | -0.080 ± 0.028 | +0.080 ± 0.028 | +0.078 ± 0.027 | -0.166 ± 0.010 | +0.162 ± 0.009 | +0.163 ± 0.011 |
| | No Rnaseq, Wsi | -0.279 ± 0.035 | +0.270 ± 0.031 | +0.284 ± 0.037 | -0.149 ± 0.010 | +0.136 ± 0.008 | +0.145 ± 0.007 |
| | No Rppa, Wsi | -0.047 ± 0.009 | +0.050 ± 0.010 | +0.056 ± 0.007 | -0.089 ± 0.008 | +0.083 ± 0.007 | +0.085 ± 0.008 |
| | No Cna, Rnaseq, Rppa | -0.125 ± 0.032 | +0.116 ± 0.033 | +0.116 ± 0.028 | -0.196 ± 0.019 | +0.181 ± 0.018 | +0.187 ± 0.018 |
| | No Cna, Rnaseq, Wsi | -0.240 ± 0.041 | +0.221 ± 0.034 | +0.242 ± 0.037 | -0.176 ± 0.017 | +0.152 ± 0.015 | +0.169 ± 0.013 |
| | No Cna, Rppa, Wsi | -0.085 ± 0.016 | +0.093 ± 0.016 | +0.091 ± 0.019 | -0.109 ± 0.010 | +0.101 ± 0.007 | +0.099 ± 0.007 |
| | No Rnaseq, Rppa, Wsi | -0.250 ± 0.049 | +0.143 ± 0.053 | +0.167 ± 0.044 | -0.172 ± 0.022 | +0.098 ± 0.027 | +0.110 ± 0.023 |
| **256** | No Cna | -0.022 ± 0.014 | +0.017 ± 0.014 | +0.017 ± 0.014 | -0.028 ± 0.007 | +0.030 ± 0.007 | +0.029 ± 0.007 |
| | No Rnaseq | -0.025 ± 0.017 | +0.026 ± 0.016 | +0.023 ± 0.016 | -0.178 ± 0.016 | +0.177 ± 0.016 | +0.177 ± 0.016 |
| | No Rppa | -0.037 ± 0.015 | +0.035 ± 0.013 | +0.035 ± 0.010 | -0.021 ± 0.006 | +0.026 ± 0.007 | +0.026 ± 0.006 |
| | No Wsi | -0.041 ± 0.015 | +0.036 ± 0.014 | +0.042 ± 0.014 | -0.056 ± 0.007 | +0.051 ± 0.005 | +0.049 ± 0.006 |
| | No Cna, Rnaseq | -0.059 ± 0.021 | +0.049 ± 0.018 | +0.055 ± 0.017 | -0.210 ± 0.021 | +0.208 ± 0.022 | +0.209 ± 0.020 |
| | No Cna, Rppa | -0.063 ± 0.012 | +0.056 ± 0.013 | +0.061 ± 0.016 | -0.049 ± 0.011 | +0.052 ± 0.010 | +0.054 ± 0.011 |
| | No Cna, Wsi | -0.057 ± 0.026 | +0.048 ± 0.018 | +0.059 ± 0.019 | -0.088 ± 0.010 | +0.087 ± 0.010 | +0.083 ± 0.011 |
| | No Rnaseq, Rppa | -0.059 ± 0.021 | +0.046 ± 0.014 | +0.050 ± 0.018 | -0.156 ± 0.015 | +0.150 ± 0.016 | +0.152 ± 0.016 |
| | No Rnaseq, Wsi | -0.285 ± 0.027 | +0.257 ± 0.022 | +0.286 ± 0.029 | -0.168 ± 0.013 | +0.152 ± 0.013 | +0.161 ± 0.014 |
| | No Rppa, Wsi | -0.058 ± 0.014 | +0.051 ± 0.007 | +0.053 ± 0.012 | -0.086 ± 0.013 | +0.083 ± 0.012 | +0.083 ± 0.011 |
| | No Cna, Rnaseq, Rppa | -0.107 ± 0.017 | +0.083 ± 0.015 | +0.091 ± 0.014 | -0.210 ± 0.020 | +0.197 ± 0.017 | +0.202 ± 0.020 |
| | No Cna, Rnaseq, Wsi | -0.329 ± 0.051 | +0.286 ± 0.048 | +0.327 ± 0.046 | -0.199 ± 0.014 | +0.175 ± 0.016 | +0.192 ± 0.016 |
| | No Cna, Rppa, Wsi | -0.110 ± 0.029 | +0.102 ± 0.023 | +0.107 ± 0.019 | -0.106 ± 0.015 | +0.106 ± 0.013 | +0.105 ± 0.014 |
| | No Rnaseq, Rppa, Wsi | -0.302 ± 0.042 | +0.178 ± 0.038 | +0.202 ± 0.042 | -0.165 ± 0.019 | +0.089 ± 0.026 | +0.111 ± 0.018 |

**Table J.** Quantitative Impact of Generative Completion across Latent Dimensions. The table compares the performance recovery for tumor stage classification and survival analysis for embedding dimensions 64, 128, and 256. Values are reported as mean ± standard deviation across 10 experimental runs.

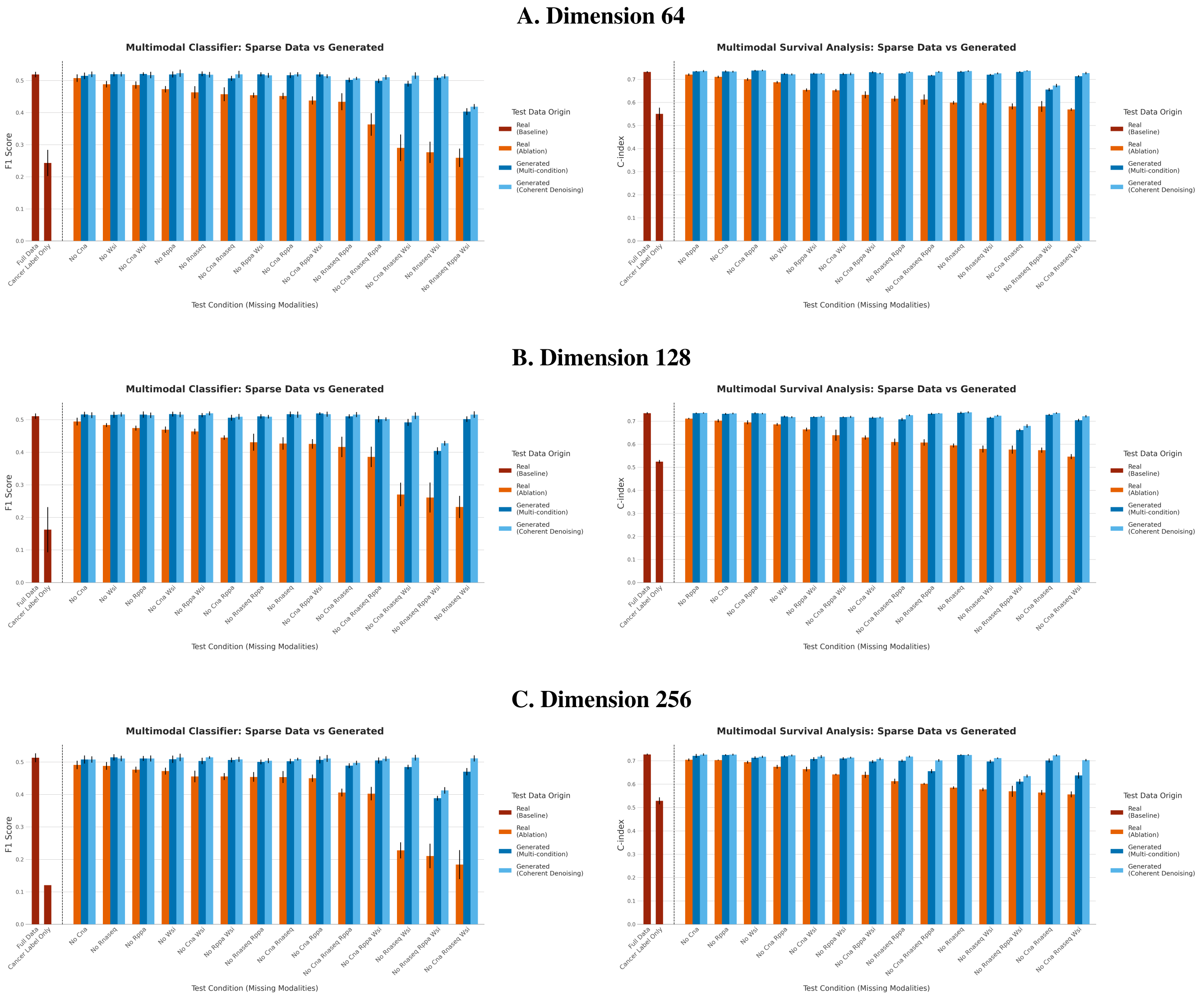


**Figure G. Downstream Performance Recovery at Higher Dimensions.** Performance comparison for tumor stage classification (Left column) and survival analysis (Right column) using embedding dimensions 64 (A), 128 (B), and 256 (C). The plots compare the Baseline (complete data) against Ablation (missing data) and the two generative imputation strategies (Multi-condition and Coherent Denoising). Error bars represent standard deviation across 10 experimental runs. The results indicate that the framework's ability to recover predictive signals remains robust and effective even as the dimensionality of the latent space increases.

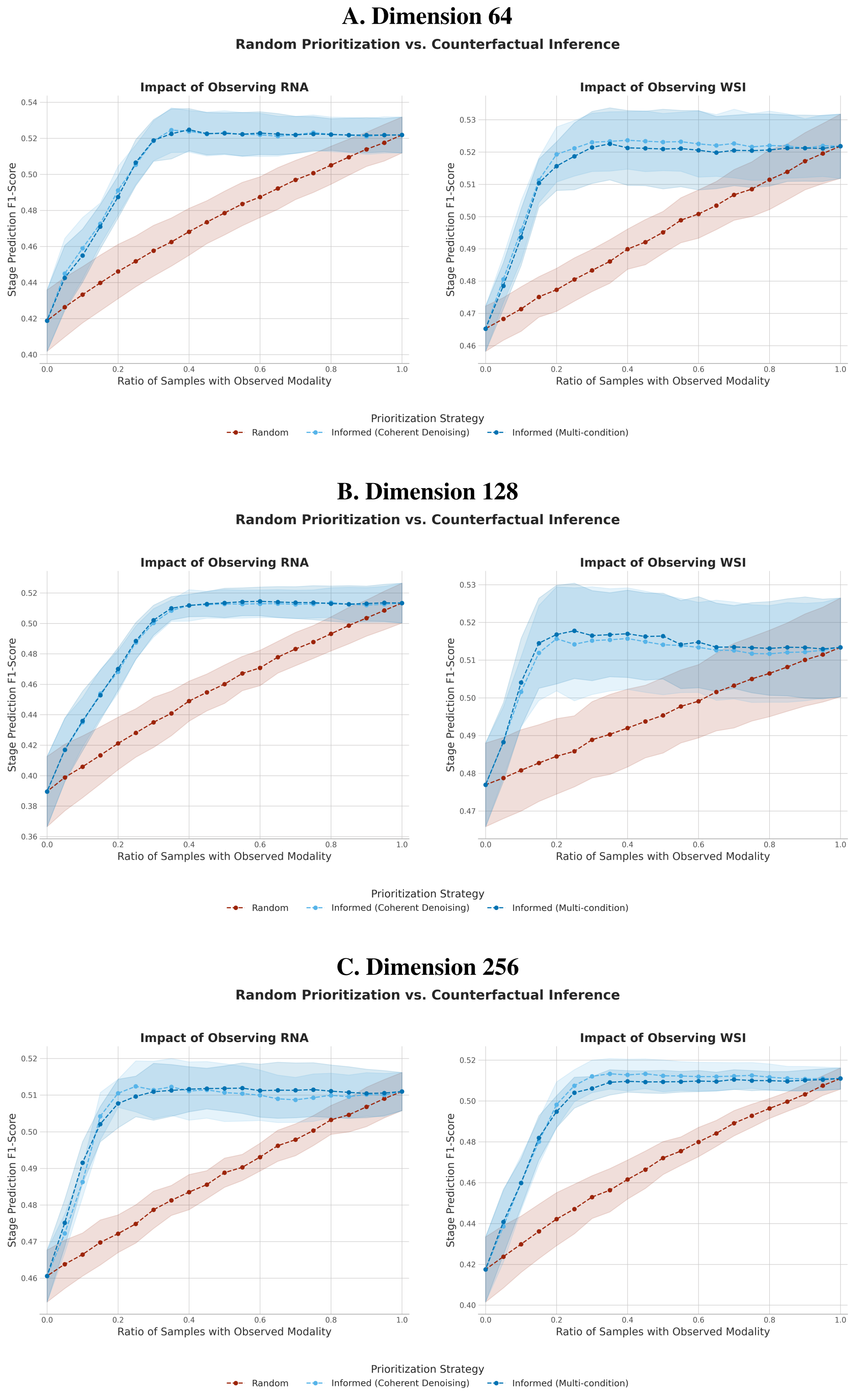


**Figure H. Robustness of Counterfactual Prioritization across Latent Dimensions.** Evaluation of the Informed Prioritization strategy for acquiring RNA-Seq (Left plots) and WSI (Right plots) data across embedding dimensions 64 (A), 128 (B), and 256 (C). The curves compare the standard Random Prioritization (red) against the model-guided Informed Prioritization (blue), which uses counterfactual variance to select the most informative patients first. Error bands indicate standard deviation across 10 repetitions. The persistent gap between the blue and red curves across all dimensions confirms that the generative model successfully captures patient-specific utility even in higher-dimensional latent spaces.
